# Favourable antibody responses to human coronaviruses in children and adolescents with autoimmune rheumatic diseases

**DOI:** 10.1101/2021.02.15.431291

**Authors:** Claire T. Deakin, Georgina H. Cornish, Kevin W. Ng, Nikhil Faulkner, William Bolland, Veera Panova, Joshua Hope, Annachiara Rosa, Ruth Harvey, Saira Hussain, Christopher Earl, Bethany R. Jebson, Meredyth G.Ll. Wilkinson, Lucy R. Marshall, Kathryn O’Brien, Elizabeth C. Rosser, Anna Radziszewska, Hannah Peckham, Judith Heaney, Hannah Rickman, Stavroula Paraskevopoulou, Catherine F. Houlihan, Moira J. Spyer, Steve J. Gamblin, John McCauley, Eleni Nastouli, Peter Cherepanov, Coziana Ciurtin, Lucy R. Wedderburn, George Kassiotis

**Affiliations:** Retroviral Immunology, The Francis Crick Institute, 1 Midland Road, London NW1 1AT, UK; Chromatin Structure and Mobile DNA Laboratory, The Francis Crick Institute, 1 Midland Road, London NW1 1AT, UK; Worldwide Influenza Centre, The Francis Crick Institute, 1 Midland Road, London NW1 1AT, UK; Signalling and Structural Biology Laboratory, The Francis Crick Institute, 1 Midland Road, London NW1 1AT, UK; Structural Biology of Disease Processes Laboratory, The Francis Crick Institute, 1 Midland Road, London NW1 1AT, UK; Centre for Adolescent Rheumatology Versus Arthritis at University College London (UCL), University College London Hospitals (UCLH), Great Ormond Street Hospital (GOSH), UCL, London, UK; UCL Great Ormond Street Institute for Child Health (ICH), UCL, London, UK; Centre for Rheumatology Research, Division of Medicine, UCL, London, UK; UCLH NHS Trust, London NW1 2BU, UK; Division of Infection and Immunity, UCL, London WC1E 6BT, UK; Department of Population, Policy and Practice, Great Ormond Street ICH, UCL, London WC1N 1EH, UK; National Institute for Health Research (NIHR) Biomedical Research Centre at GOSH, London, UK; OPAL Rheumatology Ltd, Sydney, Australia; Department of Infectious Disease, St Mary’s Hospital, Imperial College London, London W2 1NY, UK

## Abstract

Differences in humoral immunity to coronaviruses, including severe acute respiratory syndrome coronavirus 2 (SARS-CoV-2), between children and adults remain unexplained and the impact of underlying immune dysfunction or suppression unknown. Here, we examined the antibody immune competence of children and adolescents with prevalent inflammatory rheumatic diseases, juvenile idiopathic arthritis (JIA), juvenile dermatomyositis (JDM) and juvenile systemic lupus erythematosus (JSLE), against the seasonal human coronavirus (HCoV)-OC43 that frequently infects this age group. Despite immune dysfunction and immunosuppressive treatment, JIA, JDM and JSLE patients mounted comparable or stronger responses than healthier peers, dominated by IgG antibodies to HCoV-OC43 spike, and harboured IgG antibodies that cross-reacted with SARS-CoV-2 spike. In contrast, responses to HCoV-OC43 and SARS-CoV-2 nucleoproteins exhibited delayed age-dependent class-switching and were not elevated in JIA, JDM and JSLE patients, arguing against increased exposure. Consequently, autoimmune rheumatic diseases and their treatment were associated with a favourable ratio of spike to nucleoprotein antibodies.

## Introduction

Four types of human coronaviruses (HCoV) are endemic in the human population, causing frequent infection with relatively mild disease (1–4). The multiple introduction of zoonotic coronaviruses in the last couple of decades has highlighted their considerable pathogenic potential (5). This is exemplified by the pandemic caused by severe acute respiratory syndrome coronavirus 2 (SARS-CoV-2) that currently continues to spread globally (6).

The outcome of SARS-CoV-2 infection is highly variable both in presentation and prevalence. Several subtypes of coronavirus disease-19 (COVID-19) are now recognised, ranging from a mild flu-like, severe gastrointestinal or respiratory disease to multi-organ failure (6, 7). Severe complications of COVID-19 are comparatively rare and depend on age, gender, ethnicity, access to healthcare, socioeconomic status and underlying health conditions (6, 7). A sizable proportion of SARS-CoV-2 infections may also be asymptomatic, particularly in younger individuals (8). Children in particular appear relatively protected from severe COVID-19 (9). This observation mirrors findings from previous epidemics caused by SARS-CoV and Middle East respiratory syndrome coronavirus (MERS-CoV), again sparing children, although transmission of earlier zoonotic coronaviruses was much less widespread or documented (10). In contrast, children experience more frequent infections than adults with one or more of the four seasonal HCoVs (3, 11), and are more likely to harbour pre-existing antibodies and memory B cells that cross-react with SARS-CoV-2 (12, 13).

Whilst protected from severe COVID-19, unexplained inflammation following SARS-CoV-2 infection is being increasingly recognised in a very small fraction of children and adolescents (14). The causes of this condition, termed paediatric inflammatory multisystem syndrome temporally associated with SARS-CoV-2 infection (PIMS-TS) or multisystem inflammatory syndrome in children and adolescents (MIS-C) are poorly understood, but are thought to involve a dysregulated antibody response (15).

Management of the pandemic has necessitated public health measures, including shielding, aiming to protect vulnerable populations. Initial assessment in adult patients suggested increased risk of COVID-19 associated with rheumatic and other autoimmune diseases, attributable to their immunosuppressive treatment (16, 17). Children and adolescents with the most prevalent inflammatory rheumatic diseases, such juvenile idiopathic arthritis (JIA), juvenile dermatomyositis (JDM) and juvenile systemic lupus erythematosus (JSLE) were initially considered to be potentially at risk, owing to humoral immune dysfunction and associated immunosuppressive treatments. However, data on the ability of such patients to mount a humoral response to coronavirus infections is lacking.

Here, we compared the antibody responses of paediatric and adolescent JIA, JDM and JSLE patients to two immunodominant coronaviral antigens with those of their healthier peers. SARS-CoV-2 infections are relatively rare in children and adolescents with rheumatic diseases due to shielding, and relatively undetected in healthier children and adolescents due to lack of severe symptoms and limited mass testing. We, therefore, studied antibody responses to one of the seasonal HCoVs, HCoV-OC43, which causes very frequent infection in this age group (3, 11) and may also elicit antibodies that cross-react with SARS-CoV-2 (12, 18), as a surrogate of antibody response to coronaviruses in general. Although the future of the current pandemic is difficult to accurately project, recent studies suggest SARS-CoV-2 may ultimately become endemic with a low infection fatality ratio, similar to seasonal HCoVs (19). Interestingly, analysis of the mutational history of HCoV-OC43 indicated relatively recent zoonotic introduction into humans (20), which may have resulted in a major epidemic before HCoV-OC43 became endemic. Therefore, immune responses to current seasonal HCoVs can be informative of the response to SARS-CoV-2, should the latter also become endemic. Our findings suggest that inflammatory rheumatic diseases do not impede humoral immunity to a seasonal HCoV in this age group and may even enhance it.

## Results

### Normal or stronger IgG responses to coronaviral spikes in paediatric and adolescent JIA, JDM and JSLE patients

To detect and quantify antibodies reactive with HCoV-OC43 and SARS-CoV-2 spikes, we used a flow cytometry-based assay, which relies on expression of the full-length spikes on the surface of HEK293T cells (**Fig. S1**). Using this assay, we previously described the presence of SARS-CoV-2 spike-reactive antibodies in pre-COVID-19 sera collected from children and adolescents, as well as a smaller fraction of adults (12). To quantify relative antibody titres, we modified the assay by mixing equal ratios of HEK293T cells transfected to express each coronaviral spike (HEK293T.spike) with HEK293T cells transduced to express an unrelated retroviral envelope glycoprotein (HEK293T.env), which served as control, and GFP (**Fig. S1**). Cells expressing HCoV-OC43 spike were mixed with cells expressing the human endogenous retrovirus (HERV) ERV3-1 envelope glycoprotein and cells expressing SARS-CoV-2 spike were mixed with cells expressing HERV-K113 envelope glycoprotein. Specific increase in staining intensity was calculated by comparing the mean fluorescence intensity (MFI) of HEK293T.spike cells and HEK293T.env with the MFI of unstained cells.

In all healthy control and patient groups of children and young people (**Table S1**), IgG was the main class of spike-binding antibodies, representing 68% and 61% of all antibodies to HCoV-OC43 and SARS-CoV-2 spikes, respectively (**Fig. 1A**). IgM antibodies represented 27% and 30% of antibodies to HCoV-OC43 and SARS-CoV-2 spikes, respectively, which was likely an overestimate, owing to the higher non-specific staining of IgM antibodies. In contrast, IgA antibodies were the least abundant, likely due to competition with IgG antibodies in the assay, as previously observed (12).

**Figure 1.**
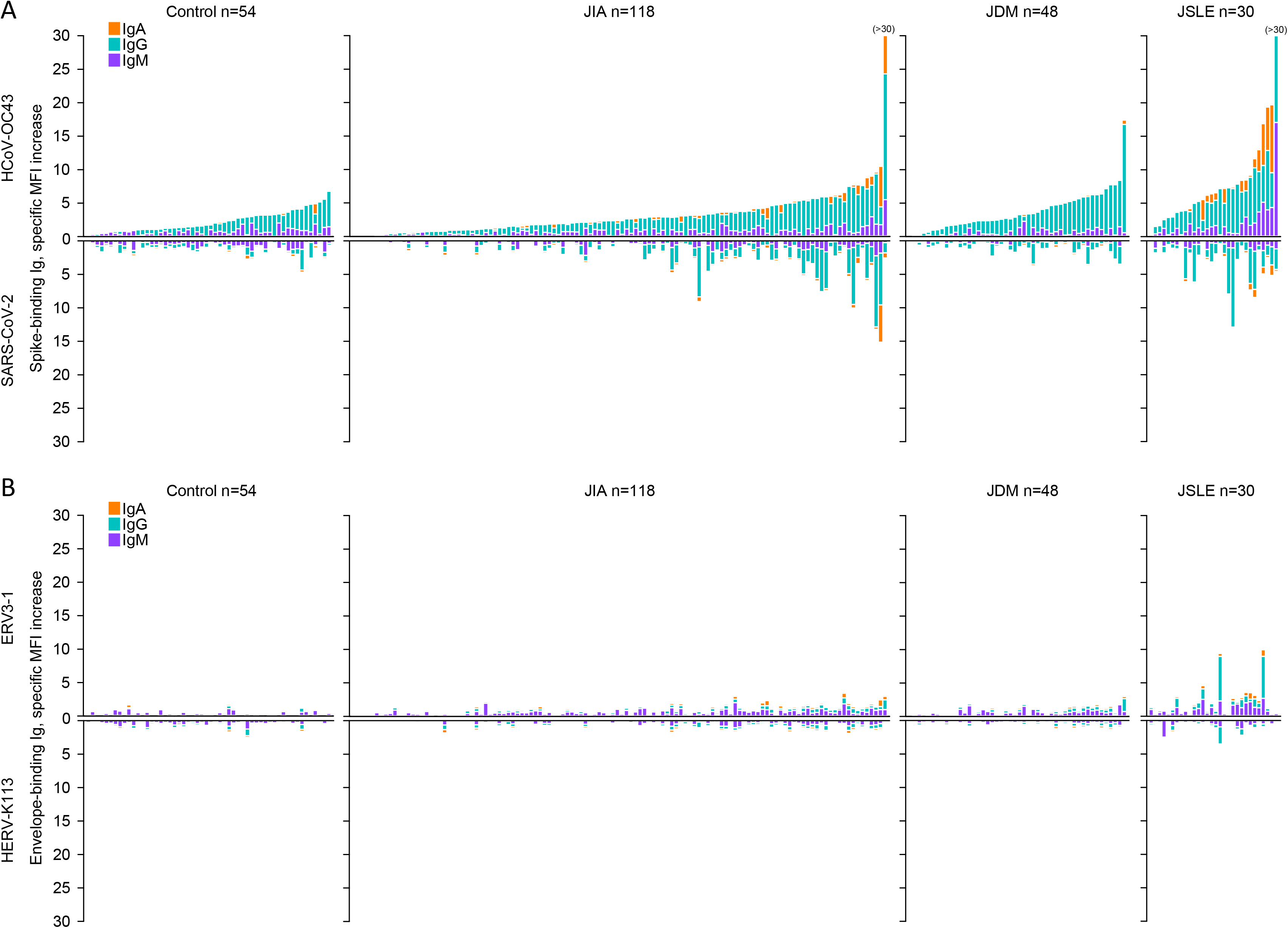
Antibodies to coronaviral spikes in paediatric and adolescent JIA, JDM and JSLE patients and age-matched controls. (A) Mirror plots of the specific MFI increase of HEK293T cells expressing HCoV-OC43 spike (top) or SARS-CoV-2 spike (bottom) caused by individual sera. Each bar is an individual healthy control or patient. Samples are plotted according to the signal of antibodies to HCoV-OC43 spike and in the same position in the mirror plots. (B) Mirror plots of the specific MFI increase of HEK293T cells expressing ERV3-1 (top) or HERV-K113 (bottom) envelope glycoproteins caused by individual sera. Samples are plotted in the same order as in A.

Consistent with our previous findings (12), the majority of healthy children and adolescents had detectable IgG antibodies to HCoV-OC43 spike, and a smaller fraction had detectable antibodies to SARS-CoV-2 spike and at much lower levels (**Fig. 1A, Table 1**). Paediatric and adolescent JIA, JDM and JSLE patients showed a similar profile, the majority having IgG antibodies to HCoV-OC43 spike and a fraction having IgG antibodies to SARS-CoV-2 spike (**Fig. 1A, Table 1**). Prevalence of IgG antibodies to HCoV-OC43 spike was significantly higher in JDM and JSLE patients than in healthy controls, as was the prevalence of IgG antibodies to SARS-CoV-2 spike in JSLE patients (**Table 1**). In contrast, antibodies to ERV3-1 or HERV-K113 envelopes were very rarely detected in any group, with the exception of JSLE, which included several adolescent patients with high IgG antibody levels to ERV3-1, two of whom also had detectable antibodies to HERV-K113 (**Fig. 1B**).

**Table 1.** Prevalence of IgG antibodies to OC43 and SARS-CoV-2 spikes in JIA, JDM and JSLE patients.

| Disease group | Prevalence of IgG to OC43 spike |  | Prevalence of IgG to SARS-CoV-2 spike |  |
| --- | --- | --- | --- | --- |
|  | positive/total (%) | *p* value<sup>1</sup> | positive/total (%) | *p* value<sup>1</sup> |
| Control | 40/54 (74.1) | – | 21/54 (38.9) | – |
| JIA | 100/118 (84.7) | ns | 57/118 (48.3) | ns |
| JDM<sup>2</sup> | 44/48 (91.7) | 0.0353 | 23/48 (47.9) | ns |
| JSLE | 29/30 (96.7) | 0.0149 | 25/30 (83.3) | 0.0001 |

<sup>1</sup>Fisher's exact tests between each disease group and the healthy control. <sup>2</sup>Values were not available for one of the JDM patients.

Given that adolescent JSLE patients were, on average, older than other patients and healthy controls (**Table S1**) and that age may influence antibody levels and class, we stratified each condition by age. We considered two age groups, 1-12 and 13-18 years of age, referred to here as younger and older, respectively. Compared with the respective healthy control, younger JIA and JDM patients had significantly higher levels of IgG antibodies to HCoV-OC43 spike, which were largely normalised in older JIA and JDM patients (**Fig. 2A**). Significantly higher levels of IgG antibodies to HCoV-OC43 spike were also observed in JSLE patients (**Fig. 2A**). As IgG antibodies were the predominant class, these differences were also reflected in the total antibodies to HCoV-OC43 spike (**Fig. 2A**). IgA antibodies to HCoV-OC43 spike were also elevated in JSLE and older JIA patients (**Fig. 2A**). Despite the fact that antibodies to SARS-CoV-2 spike were found at much lower levels and correlated weakly with antibodies to HCoV-OC43 spike (correlation coefficient 0.376 for IgG antibodies, p<0.0001), their prevalence in paediatric and adolescent JIA, JDM and JSLE patients mirrored antibodies to HCoV-OC43 spike (**Fig. 2B**). Significantly higher levels of IgG antibodies to SARS-CoV-2 spike were observed in JSLE and younger JIA patients, whereas, total antibodies to SARS-CoV-2 spike seemed to drop below control levels in older JIA patients (**Fig. 2B**).

**Figure 2.**
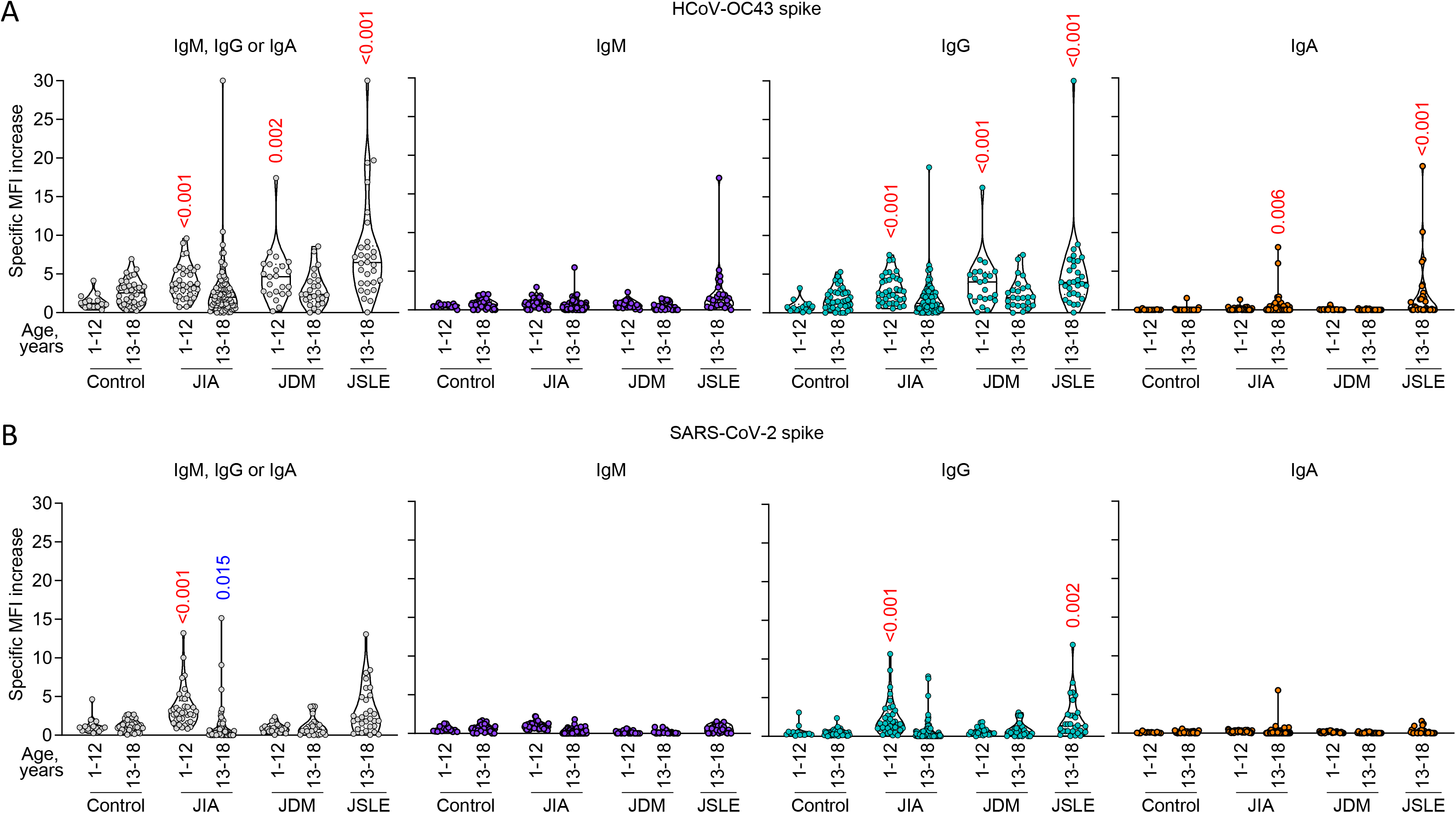
Antibodies to coronaviral spikes in paediatric and adolescent JIA, JDM and JSLE patients and controls of different age. Specific MFI increase of HEK293T cells expressing HCoV-OC43 spike (A) or SARS-CoV-2 spike (B) caused by individual sera from the indicated age and disease group. Each symbol is an individual healthy control or patient. Red and blue numbers within the plots denote the p values of statistically significant increases and decreases, respectively, when comparing each disease group with the respective healthy control of the same age group. The older control group was also compared with the younger control group.

In addition to the effect of the disease group, multiple regression analysis suggested an independent effect of multiple other factors, including donor or patient gender, age, year of sampling, immunosuppressive treatment and autoantibody presence. After controlling for the effects of female sex, older age, year of sample collection and steroid treatment, total antibodies to HCoV-OC43 spike were significantly higher in JSLE patients compared to controls, while IgG antibodies to HCoV-OC43 spike were elevated in paediatric and adolescent patients of all three conditions (**Tables S2-S3**). Female sex and steroid treatment at the time of sampling were associated with higher antibody levels. JDM was associated with lower levels of total antibodies to SARS-CoV-2 spike while JSLE was associated with higher levels of IgG antibodies to SARS-CoV-2 spike, after the effects of female sex, older age, year of sample collection and autoantibody presence were controlled for. Total and IgG antibodies to SARS-CoV-2 spike were elevated in younger patients and in those with autoantibodies (**Tables S4-S5**).

As these rheumatic diseases affect more females than males, the former are overrepresented in the JIA, JDM and JSLE groups (**Table S1**). Nevertheless, when only younger male patients were compared with younger male healthy controls, levels of IgG antibodies to HCoV-OC43 spike were still significantly higher in JIA and JDM (p=0.00000993 and p=0.000728, respectively, Student’s t-test), demonstrating that this effect is independent of patient gender.

The effect of the year of sampling may reflect the variation in HCoV-OC43 prevalence between winters in the UK (21), affecting the strength of the contemporaneous antibody response. Alternatively, it may reflect antigenic drift in HCoV-OC43 over long periods of time, leading to escape from antibody response, as recently suggested for HCoV-229E (22). HCoV-OC43 spike sequence we used in this study was from a clone isolated in 2017 and it was, therefore, possible that it was better recognised by more recent, than older sera (22). However, no linear relationship was observed between titres of antibodies to HCoV-OC43 spike and sampling year (**Fig. S2**), arguing against the possibility that the 2017 HCoV-OC43 spike was not recognised by older sera. A similar pattern was seen also for antibodies that cross-reacted with SARS-CoV-2 spike (**Fig. S2**), which are likely targeting more conserved regions of the spike and would not be affected by escape mutations in HCoV-OC43 spike.

The positive association between immunosuppressive steroid treatment and antibodies to coronaviral spike proteins was unexpected and remained even when other factors were taken into account (**Tables S2-S3**). Although it may not simply be a surrogate for disease activity, necessitating higher doses, steroid treatment may still capture other underlying aspects of immune dysregulation, particularly in the JSLE group.

Lastly, the positive correlation between antibodies to HCoV-OC43 and SARS-CoV-2 spikes and autoantibodies (e.g. antinuclear antibodies detected as part of diagnosis) was additionally supported by similar correlation also with antibodies to HERV-encoded envelope glycoproteins, which are effectively autoantibodies. Indeed, we observed a correlation between IgG to HCoV-OC43 or SARS-CoV-2 spikes and IgG autoantibodies to ERV3-1 (correlation coefficients 0.692, p<0.0001; and 0.457, p<0.0001; respectively) or IgG autoantibodies to HERVK-113 (correlation coefficients 0.277, p<0.0001; and 0.681, p<0.0001; respectively), indicating a common underlying cause, such as a hyperactive adaptive immune system.

As an independent method of antibody detection and also of potential functional relevance, we also assessed the ability of sera to neutralise SARS-CoV-2 *in vitro*. Most, but not all of the sera with SARS-CoV-2 S-cross-reactive IgG, detected by flow cytometry, also exhibited *in vitro* SARS-CoV-2 neutralising activity (**Fig. 3A**). SARS-CoV-2 neutralisation titres were similar between the three conditions and between age groups, and comparable with those previously reported in healthy children and adolescents (**Fig. 3A**). Neutralisation activity showed a weak correlation with IgG to SARS-CoV-2 spike, which was expected given that not all spike-binding antibodies are neutralising, but no correlation with IgG to HCoV-OC43 spike (**Fig. 3B**).

**Figure 3.**
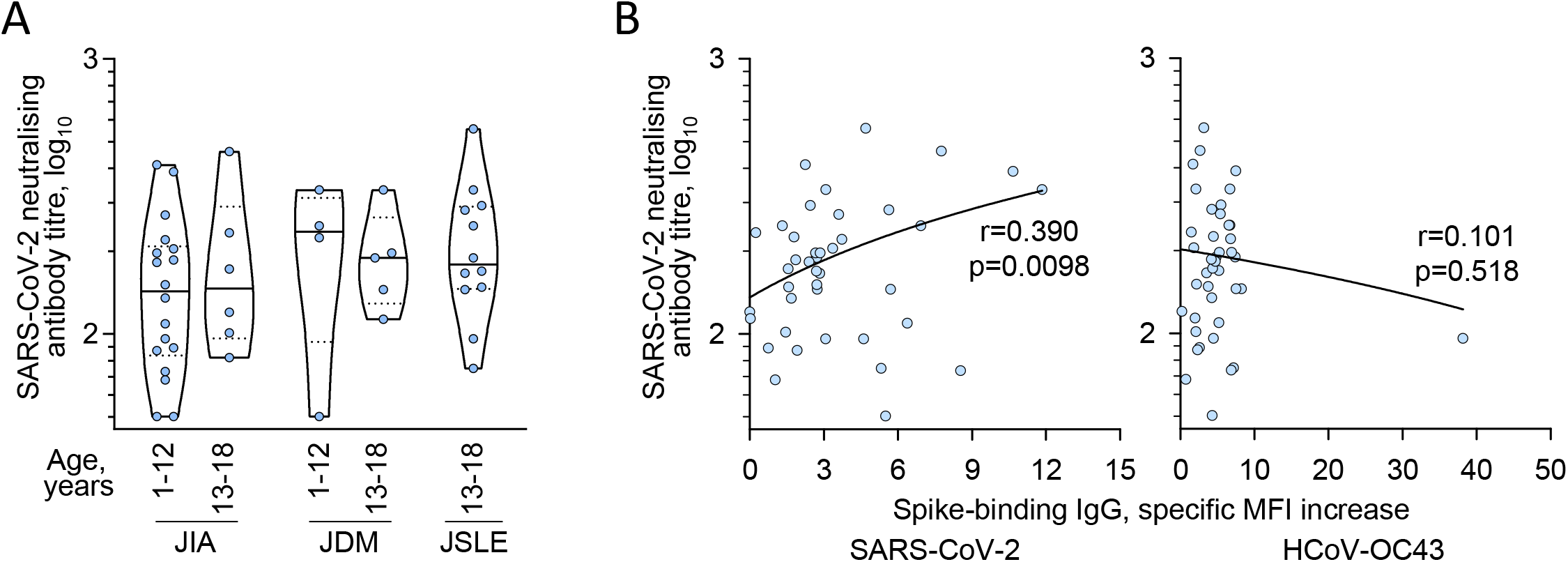
SARS-CoV-2 neutralising antibodies in paediatric and adolescent JIA, JDM and JSLE patients. (A) SARS-CoV-2-neutralising antibody titres in the indicated age and disease group. Only patients with SARS-CoV-2 spike-binding antibodies detectable by flow cytometry were included. (B) Correlation of SARS-CoV-2-neutralising antibody titres with levels of flow cytometry-detectable antibodies to SARS-CoV-2 spike (left) or HCoV-OC43 spike (right). In A and B, each symbol is an individual patient.

Collectively, these data suggest that paediatric and adolescent JIA, JDM and JSLE patients mounted a comparable, if not stronger class-switched response to HCoV-OC43 spike, compared with healthy children and adolescents, and harboured cross-reactive antibodies to SARS-CoV-2 spike.

### Predominant IgM responses to coronaviral nucleoproteins in children and adolescents

Stronger IgG responses to coronaviral spikes in paediatric and adolescent JIA, JDM and JSLE patients than in healthy age-matched controls could potentially arise from immune hyperactivity to comparable exposure to HCoVs, as may be expected from the underlying rheumatic disease. Alternatively, it could arise from higher exposure to or infection with HCoVs in these patients, potentially due to immunosuppressive treatments. To explore these two possibilities, we assessed responses to a second coronaviral antigen, the nucleoprotein. As their target is an internal viral protein, antibodies to nucleoproteins are not neutralising and their levels are thought to better reflect the amount of viral replication than antibodies to spikes.

To quantify antibodies to HCoV-OC43 and SARS-CoV-2 nucleoproteins, we developed a quantitative bead-based assay (**Fig. S3**). Using this assay, we were able to detect HCoV-OC43 nucleoprotein-binding antibodies in effectively all samples (99.6%) and SARS-CoV-2 nucleoprotein-binding antibodies in a substantial proportion (70.1%) (**Fig. 4A**). Surprisingly, however, the dominant class of nucleoprotein-binding antibodies in these cohorts was IgM, making up to 82.2% and 67.3% of antibodies to HCoV-OC43 and SARS-CoV-2 nucleoproteins, respectively. This finding contrasted with the dominance of IgG antibodies to coronaviral spikes in children found here and in prior reports (23), and also with reports of class use in antibodies to other antigens in children versus adults in general (13). We therefore examined a group of adults, across the age spectrum, to establish whether the apparent dominance of IgM antibodies to nucleoproteins were due the assay used or indeed a unique feature of the response of children and adolescents to these nucleoproteins in particular. Healthy adults responded to HCoV-OC43 nucleoprotein with a balanced IgG and IgM response, comprising 31.4% and 61.9% of the total response to this antigen. (**Fig. 4B**).

**Figure 4.**
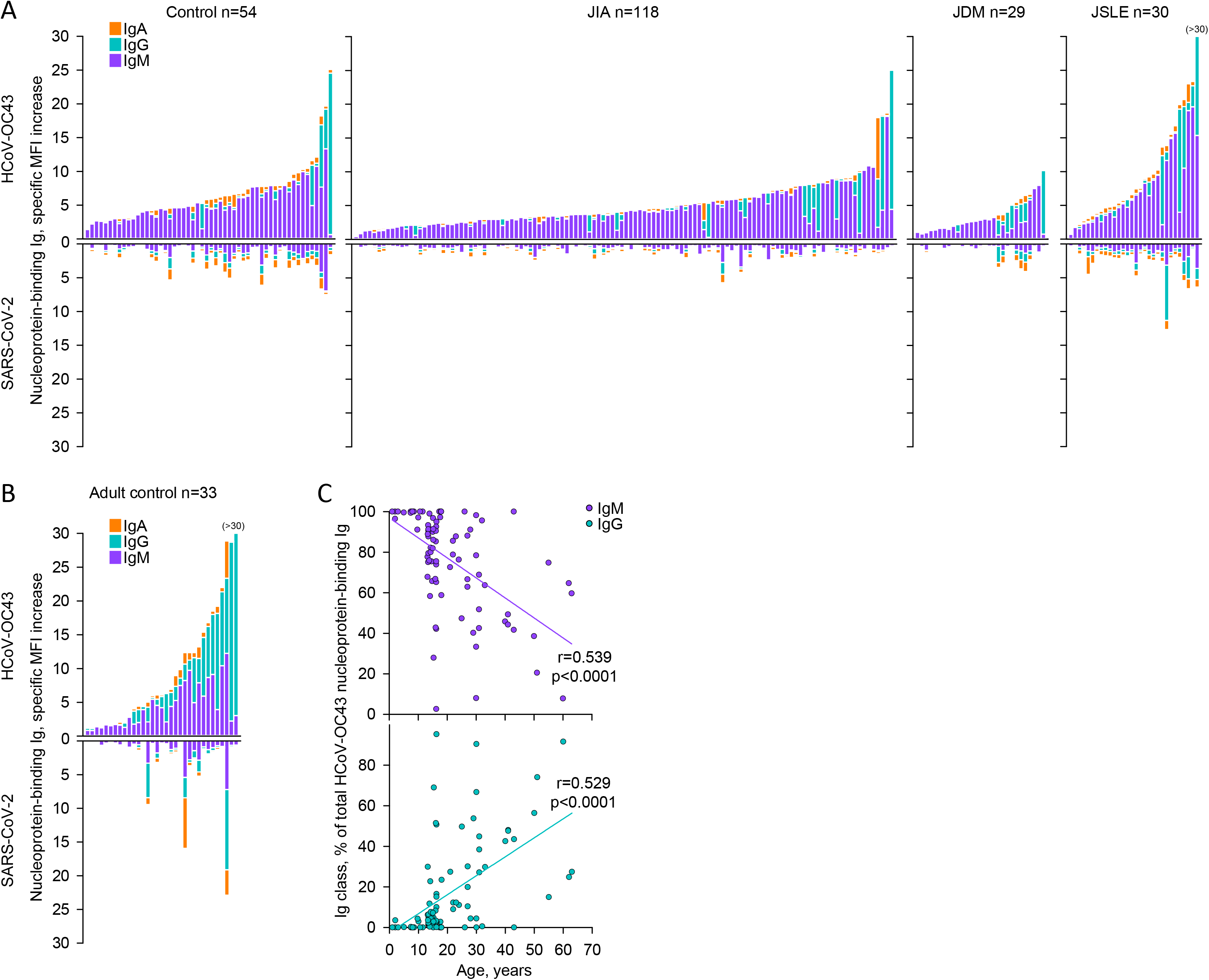
Antibodies to coronaviral nucleoproteins in paediatric and adolescent JIA, JDM and JSLE patients and age-matched controls. Mirror plots of the specific MFI increase of HEK293T cells expressing HCoV-OC43 nucleoprotein (top) or SARS-CoV-2 nucleoprotein (bottom) caused by individual sera. Each bar is an individual child or adolescent healthy control or patient (A) or adult healthy control (B). Samples are plotted according to the signal of antibodies to HCoV-OC43 nucleoprotein and in the same position in the mirror plots. (C) Correlation of the proportion of total HCoV-OC43 nucleoprotein-binding antibodies represented by IgM (top) or IgG (bottom) classes, and the age of the donor or patient. Each symbol is an individual sample.

This response of adults was significantly skewed compared with that of healthy children and adolescents, which comprised 8.7% IgG and 84.6% IgM (p<0.001 when comparing the IgG or IgM titres to HCoV-OC43 nucleoprotein between healthy children/adolescents and healthy adults, Mann-Whitney U test). Thus, children and adolescents made an antibody response to coronaviral nucleoproteins that were unusually dominated by IgM and that gradually and partially switched to IgG with age (**Fig. 4C**).

In comparison with the respective healthy control group, paediatric and adolescent JIA, JDM and JSLE patients showed no evidence for elevated antibody responses to HCoV-OC43 nucleoprotein (**Fig. 5A**). These responses were in fact reduced in certain groups such as older JIA and younger JDM patients (**Fig. 5A**). A similar profile was observed also for antibodies to SARS-CoV-2 nucleoprotein, with a significant increase with age in the healthy control group, and decrease in older JIA patients compared with their healthy control (**Fig. 5B**). Multiple regression models confirmed that total antibodies and IgM antibodies to both HCoV-OC43 and SARS-CoV-2 nucleoproteins were reduced in JDM and JIA patients, after controlling for the effects of female sex and older age (**Tables S6-S9**).

**Figure 5.**
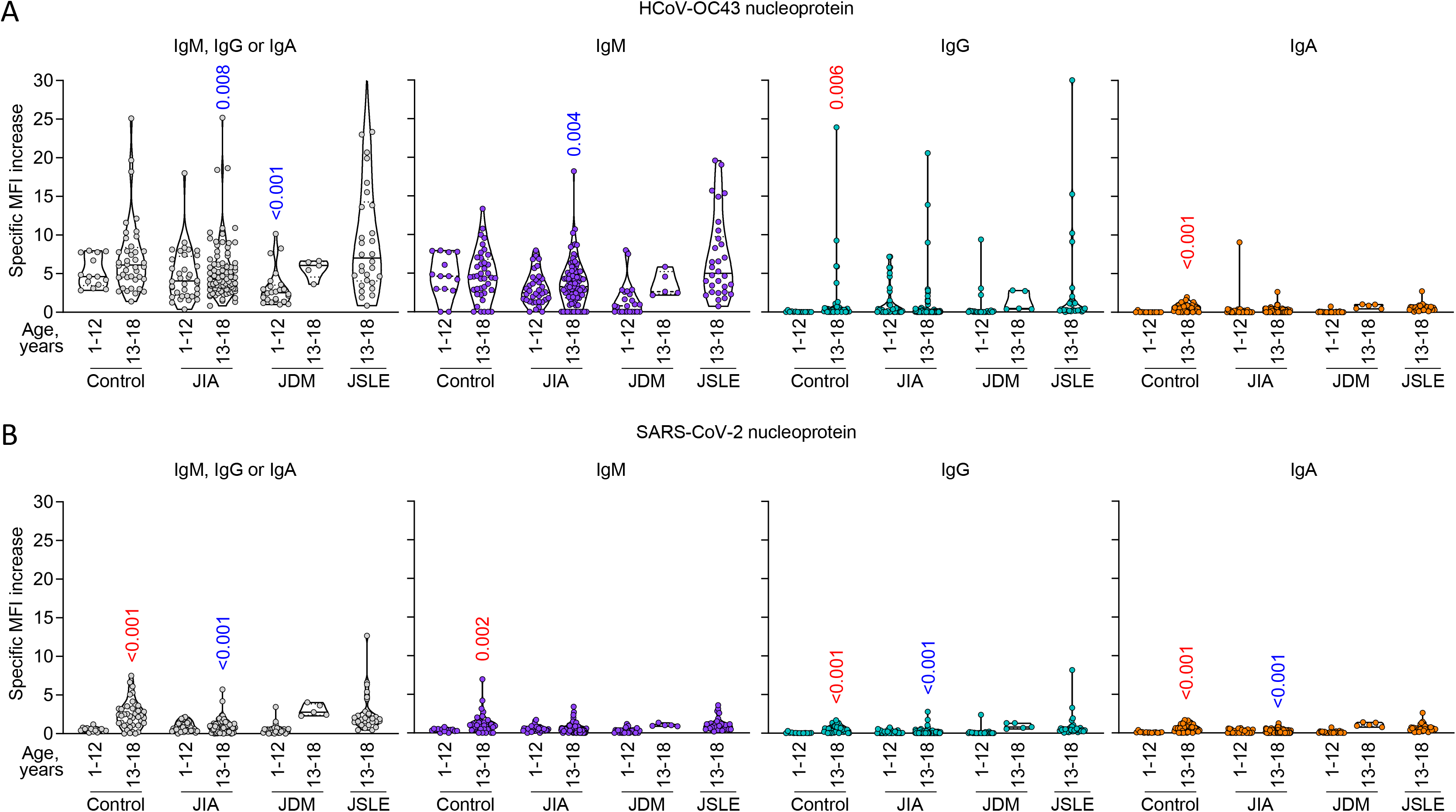
Antibodies to coronaviral nucleoproteins in paediatric and adolescent JIA, JDM and JSLE patients and controls of different age. Specific MFI increase of HEK293T cells expressing HCoV-OC43 nucleoprotein (A) or SARS-CoV-2 nucleoprotein (B) caused by individual sera from the indicated age and disease group. Each symbol is an individual healthy control or patient. Red and blue numbers within the plots denote the p values of statistically significant increases and decreases, respectively, when comparing each disease group with the respective healthy control of the same age group. The older control group was also compared with the younger control group.

These experiments revealed that children and adolescents respond differently to coronaviral spikes and nucleoproteins, in terms of magnitude and antibody class. Moreover, across all paediatric and adolescent groups, the antibody responses to the spike and nucleoprotein of each virus was only weakly correlated (correlation coefficients 0.241 and 0.286, for HCoV-OC43 and SARS-CoV-2, respectively), suggesting, differential targeting of either antigen in a given patient or donor. Preferential targeting of SARS-CoV-2 spike rather than nucleoprotein has been linked with favourable outcome (24, 25), and we therefore examined the ratio of antibodies to spike and nucleoprotein in the patient cohorts. This analysis revealed significantly higher ratios of HCoV-OC43 spike to nucleoprotein antibodies in all disease groups, compared with the respective healthy control group, with the exception of older JIA patients (**Fig. 6**). Higher ratios of spike to nucleoprotein antibodies for all disease groups and for both HCoV-OC43 and SARS-CoV-2 was confirmed by multiple regression analysis, which also identified significant positive associations with female sex and steroid treatment, and a negative association with older age (**Tables S10-S11**). This profile was consistent with lower exposure and a better outcome of infection in these patients.

**Figure 6.**
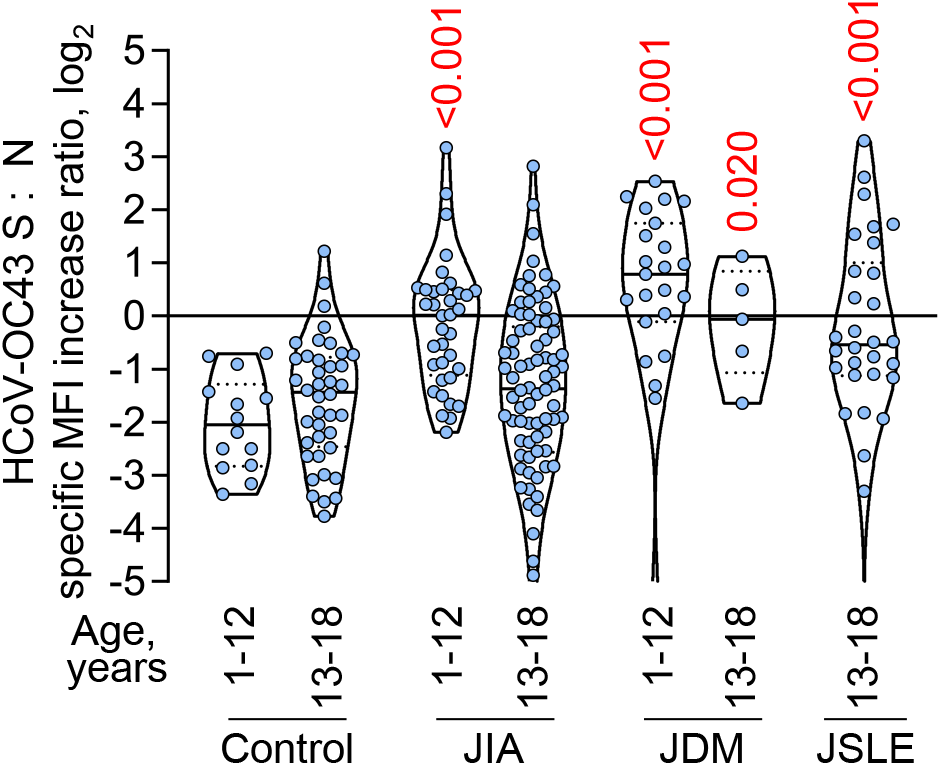
Ratio of levels of antibody to coronaviral spikes and nucleoproteins in paediatric and adolescent JIA, JDM and JSLE patients and age-matched controls. The log2-transformed ratios of total antibodies to HCoV-OC43 spike to total antibodies to HCoV-OC43 nucleoprotein (S : N) are plotted for the indicated age and disease group. Each symbol is an individual sample. Numbers within the plots denote the p values of statistically significant increases, when comparing each disease group with the respective healthy control of the same age group.

## Discussion

The current pandemic caused by SARS-CoV-2 infection has highlighted the need for deeper understanding of the interaction of coronaviruses with the human immune system. Antibody responses, as well as the outcome of SARS-CoV-2 infection, are highly variable among healthy adults, for reasons that remain incompletely understood (26), and may be even more variable in people with immunological disorders. Here, we show that, despite adaptive immune dysfunction and immunosuppressive treatment, paediatric and adolescent JIA, JDM and JSLE patients mount antibody responses to coronaviruses that are comparable to or stronger than those of healthier peers.

The susceptibility of patients with common rheumatic or other autoimmune diseases to SARS-CoV-2 infection and subsequent disease is still a matter of debate (27). Studies conducted exclusively in adult cohorts suggested an increased risk of hospitalisation due to COVID-19 in autoimmune disease patients, which was attributed to immunosuppressive treatment with glucocorticoids particularly in rheumatic diseases (16, 17, 28), an association seemingly at odds with the observed effect of dexamethasone treatment in lowering mortality in patients with the most severe COVID-19 symptoms (29). Moreover, independent studies indicated that increased risk of COVID-19 in patients with rheumatic diseases may not be associated with immunosuppressive treatment directly, but rather with disease activity, which requires proportionally higher doses of glucocorticoids (30). Lastly, even the association between increased risk of COVID-19 and rheumatic disease may not be direct. A study of rheumatic disease patients in a region with high COVID-19 prevalence linked the risk of COVID-19 with age and comorbidities, instead of type of rheumatic condition or immunosuppressive treatment (31).

Rheumatic diseases include conditions with varying aetiology and degree of immune dysfunction, which likely contributes to the complexity of association with the risk of COVID-19. Nevertheless, the severity of COVID-19 in this group of patients appears much less than was originally expected. Moreover, children are at a much lower risk of COVID-19 than adults in general, and studies of COVID-19 susceptibility in children with rheumatic diseases are currently lacking.

Using the highly prevalent HCoV-OC43, which frequently infects children (3, 11), we found no evidence for a defective antibody response to the spike glycoprotein in JIA, JDM and JSLE patients. In fact, all other variables considered, such patients often mounted a stronger response to the HCoV-OC43 spike, likely due to immune hyperactivity, despite immunosuppressive treatment. Consistent with these findings, pre-existing antibodies that cross-react with SARS-CoV-2 spike and cross-neutralise authentic SARS-CoV-2 *in vitro*, were also higher in paediatric and adolescent JIA, JDM and JSLE patients, arguing against deficient immunity to coronaviruses. Our findings are consistent with a study of SARS-CoV-2 infection in a small number of adult rheumatic diseases patients, which detected antibody responses in a majority (32).

Levels of spike-reactive antibodies were also affected to a certain degree by the year in which the samples were collected. A recent report provided evidence to suggest antigenic drift of HCoVs, likely driven by antibody-mediated selection (22). Indeed, antibodies in historical human serum samples neutralised contemporaneous isolates of HCoV-229E, but not future isolates (22). The relationship between collection year and HCoV-OC43 spike-reactive antibody levels we observed here was not linear, with no overall change over time. This finding does not speak against potential antigenic drift (22) for the following reasons. Firstly, the time interval we analysed here was relatively shorter and, secondly, we measured binding antibodies, which recognise a wider range of antigenic epitopes than neutralising antibodies and, therefore, are less affected by specific mutations.

In contrast to antibodies to coronaviral spikes, those to nucleoproteins were not found elevated in paediatric and adolescent JIA, JDM and JSLE patients, and in certain groups were even reduced, compared with healthy controls. This finding argues against increased exposure to HCoVs in such patients that could have resulted from relative immune deficiency, but rather to a stronger response to the spikes following similar or possible lower exposure.

Interestingly, whereas both pre-existing and *de novo* antibody responses of children and adolescents to SARS-CoV-2 spike is predominantly of the IgG class (12, 23), we found that their responses to the nucleoproteins are dominated by IgM. This feature highlights distinct paths of evolution of the antibody responses to spike and nucleoprotein, with much slower and incomplete class switch for the latter with age. Of note, dominance of IgM and IgG antibodies in healthy children and elderly, respectively were also observed in the response to the nucleoprotein of another seasonal coronavirus, HCoV-229E, in a systems serology study (33), although this feature extended also to the spike in that study.

The early balance of antibodies to SARS-CoV-2 spike and nucleoprotein has been linked with COVID-19 survival, with lower ratios associated with increased risk of death in adult patients (24). In an independent study, higher ratios of antibodies to SARS-CoV-2 spike and nucleoprotein were observed in COVID-19 patients with mild illness than in severely ill patients (25). These findings supported a model where higher ratio of protective antibodies to spike and non-protective antibodies to nucleoprotein is a hallmark of effective immunity, predicting disease trajectory (24). In the case of HCoV-OC43, the ratio of antibodies to spike and nucleoprotein was significantly higher in paediatric and adolescent JIA, JDM and JSLE patients. If the response to SARS-CoV-2 or vaccination can be inferred from the response to HCoV-OC-43, together with the presence of pre-existing cross-reactive antibodies to SARS-CoV-2, the data suggest that paediatric and adolescent patients with the most common inflammatory rheumatic diseases would mount a highly effective humoral response against SARS-CoV-2 infection or vaccination.

## Methods

### Donor and patient samples and clinical data

Pre-pandemic serum or plasma samples from children and adolescents were obtained from the Centre for Adolescent Rheumatology Versus Arthritis at University College London (UCL), University College London Hospitals (UCLH), and Great Ormond Street Hospitals (GOSH), and UCL Great Ormond Street Institute for Child Health (ICH) with ethical approval (refs 11/LO/0330, 99RU11, 04RU07 and 95RU04). To balance the age range between healthy donors and patients, we included 6 adolescents with gender dysphoria in our healthy adolescent group. We considered gender dysphoria patients immunologically healthy. This group was analysed according to gender assigned at birth, to reflect chromosomal influences on adaptive immunity. Pre-pandemic samples from adults were obtained from UCLH (ref 284088). These samples were from residual samples prior to discarding, in accordance with Royal College Pathologists guidelines and the UCLH Clinical Governance for assay development and GOSH and ICH regulations. Samples had undergone at least one cycle of thaw and freeze, and had been stored at either −20°C or −80°C freezers at local hospitals prior to transfer to the Francis Crick Institute. All serum or plasma samples were heat-treated at 56°C for 30 min prior to testing.

Demographic and clinical data were obtained via the Centre for Adolescent Rheumatology Versus Arthritis, the Childhood Arthritis Response to Medication Study (CHARMS), the JIA Pathogenesis Study, the JDM Cohort & Biomarker Study databases, and medical records. Since certain items of clinical data were recorded differentially across disease cohorts or were not recorded for healthy children and adolescents, the following assumptions were applied in order to maximise complete data for the regression analyses. Patients were treated as being ‘autoantibody positive’ if any of rheumatoid factor (RF), anti-nuclear antibodies (ANA) or myositis-specific/myositis-associated autoantibodies were recorded as positive. Healthy controls were not tested for RF/ANA or myositis autoantibodies and although a low rate of low titre ANA can be detected in healthy individuals, these individuals were assumed autoantibody-negative for the purpose of this analysis. Healthy children and adolescents were assumed to not be on steroids, biologics, disease modifying anti-rheumatic drugs (DMARDs) or other immunosuppressants at time of sampling. Disease activity in patients had been recorded using either the SLE disease activity index (SLEDAI) for JSLE or a physician’s visual analogue scale (PhysVAS) for JIA and JDM. A categorical disease activity variable was defined as “no disease” (SLEDAI=0 or PhysVAS=0), “mild” (0<SLEDAI≤4 or 0<PhysVAS≤3.5) or “moderate to severe” (SLEDAI>4 or PhysVAS>3.5). Healthy controls were included in the “no disease” group. Disease duration was discretised as “no disease/diagnosis” for samples from healthy controls or from patients at diagnosis, “≤24 months” or “>24 months”, with the 24 month cut-off selected according to the distribution of disease duration.

### Cell lines and plasmids

HEK293T cells and Vero E6 were obtained from the Cell Services facility at The Francis Crick Institute and verified as mycoplasma-free. HEK293T cells were validated by DNA fingerprinting. Cells were grown in Iscove’s Modified Dulbecco’s Medium (Sigma Aldrich) supplemented with 5% fetal bovine serum (Thermo Fisher Scientific), L-glutamine (2 mM, Thermo Fisher Scientific), penicillin (100 U/ml, Thermo Fisher Scientific), and streptomycin (0.1 mg/ml, Thermo Fisher Scientific).

HEK293T cells expressing HERV-K113 envelope glycoprotein were generated by retroviral transduction with the vector encoding the putative ancestral protein sequence of HERV-K113 envelope glycoprotein (34) and GFP separated by an internal ribosome entry site (IRES). HEK293T cells expressing ERV3-1 envelope glycoprotein were similarly generated by retroviral transduction with the vector encoding the ERV3-1 envelope glycoprotein (NCBI Reference Sequence: NM_001007253.4) and GFP separated by an IRES. Transduced cells were sorted based on GFP expression to >98% purity on a FACSAria Fusion cell sorter (Beckton Dickinson) and maintained as separate cell lines.

HEK293T cells expressing SARS-CoV-2 or HCoV-OC43 spikes were generated by transient transfection, using GeneJuice (EMD Millipore), with expression plasmids, encoding each spike and were used two days after transfection. The expression vector (pcDNA3) carrying a codon-optimized gene encoding the wild-type SARS-CoV-2 spike (UniProt ID: P0DTC2) was kindly provided by Massimo Pizzato, University of Trento, Italy. The expression vector (pCMV3) carrying a codon-optimized gene (NCBI Reference Sequence: AVR40344.1) encoding the HCoV-OC43 spike of isolate HCoV-OC43/USA/TCNP_00212/2017 was obtained from SinoBiological.

### Flow cytometric detection of antibodies to spike and envelope glycoproteins

HEK293T cells expressing HCoV-OC43 spike were mixed with cells expressing ERV3-1 envelope glycoprotein and GFP and HEK293T cells expressing SARS-CoV-2 spike were mixed with cells expressing HERV-K113 envelope glycoprotein and GFP at equal rations. All cells were tryspinised prior to mixing and cell mixtures were dispensed into V-bottom 96-well plates (20,000-40,000 cells/well). Cells were incubated with sera (diluted 1:50 in PBS) for 30 min, washed with FACS buffer (PBS, 5% BSA, 0.05% sodium azide), and stained with BV421 anti-IgG (clone HP6017, Biolegend), APC anti-IgM (clone MHM-88, Biolegend) and PE anti-IgA (clone IS11-8E10, Miltenyi Biotech) for 30 min (all antibodies diluted 1:200 in FACS buffer). Cells were washed with FACS buffer and fixed for 20 min in CellFIX buffer (BD Bioscience). Samples were run on a Ze5 analyzer (Bio-Rad) running Bio-Rad Everest software v2.4 or an LSR Fortessa with a high-throughput sampler (BD Biosciences) running BD FACSDiva software v8.0, and analyzed using FlowJo v10 (Tree Star Inc.) analysis software, as previously described (12). Specific increase in staining intensity by sample antibodies was calculated by comparing the mean fluorescence intensity (MFI) of each of the four HEK293T cell types stained with serum or plasma samples for each Ig class with the MFI of the respective unstained HEK293T cell type, using the following formula: (MFI of stained cells − MFI of unstained cells)/MFI of unstained cells. Samples with values over 0.5 were considered positive for binding antibodies.

### Recombinant protein production

Full-length SARS-CoV-2 and HCoV-OC43 nucleoproteins were produced by expression in *Escherichia coli* C43(DE3) cells (Lucigen) and purified as previously described (12). The SARS-CoV-2 nucleoprotein was produced with an N-terminal His_6_ tag from pOPTH-1124 plasmid (kindly provided by Jakub Luptak and Leo James, Laboratory for Molecular Biology, Cambridge, UK). The HCoV-OC43 nucleoprotein (NCBI Reference Sequence: AOL02457.1 from isolate HCoV-OC43/human/Mex/LRTI_238/2011) was produced with an N-terminal SUMO-His6 tag, which was removed during purification, by incubation with ubiquitin-like-specific protease-1 (ULP1) at 4°C overnight. Following incubation, te protein was diluted to 300 mM NaCl in 20 mM HEPES pH 8.0, loaded onto a 5 ml HiTrap heparin column (GE healthcare) and eluted over a linear 0.3-1M NaCl gradient. Purified nucleoproteins were concentrated by ultrafiltration using appropriate VivaSpin devices (Sartorius), were snap-frozen in liquid nitrogen in small aliquots and stored at −80°C.

### Flow cytometric detection of antibodies to nucleoproteins

Aliquots of recombinant HCoV-OC43 and SARS-CoV-2 nucleoproteins were conjugated with aldehyde functionalized polymethylmethacrylate-based microspheres (PolyAn GmbH, Berlin, Germany), according to manufacturer’s instructions, and kept at 4°C until use. Transparent beads of 5 μM and 2 μM diameter were conjugated with HCoV-OC43 and SARS-CoV-2 nucleoproteins, respectively. Beads were mixed at equal ratio and were dispensed into V-bottom 96-well plates (a total of 50,000-70,000 beads/well). Beads were then stained with serum and plasma samples, as described above for HEK293T cells used in flow cytometric detection of antibodies to spike and envelope glycoproteins. Specific increase in staining intensity by sample antibodies was calculated by comparing the mean fluorescence intensity (MFI) of each type of bead stained with serum or plasma samples for each Ig class with the MFI of the respective unstained bead type, using the following formula: (MFI of stained beads − MFI of unstained beads)/MFI of unstained beads. Samples with values over 0.5 were considered positive for binding antibodies.

### SARS-CoV-2 neutralisation assay

Titres of SARS-CoV-2 neutralising antibodies were determined as previously described (12). Briefly, the SARS-CoV-2 isolate hCoV-19/England/02/2020 was obtained from the Respiratory Virus Unit, Public Health England, UK, (GISAID EpiCov™ accession EPI_ISL_407073) and propagated in Vero E6 cells. Triplicate cultures of Vero E6 cells were incubated with SARS-CoV-2 and twofold serial dilutions of human sera (previously heat-treated at 56°C for 30 min) for 3 hours. The inoculum was then removed and cells were overlaid with virus growth medium containing 1.2% Avicel (FMC BioPolymer). At 24 hours post-infection, cells were fixed in 4% paraformaldehyde and permeabilized with 0.2% Triton X-100/PBS, and virus plaques were visualized by immunostaining (12). Virus plaques were quantified and IC_50_ values were calculated using LabView software as described previously (35) or SigmaPlot v14.0 (Systat Software).

### Statistical analyses

Data were analysed and plotted in GraphPad Prism v8 (GraphPad Software) or SigmaPlot v14.0 (Systat Software). Parametric comparisons of normally-distributed values that satisfied the variance criteria were made by unpaired Student’s t-tests or One Way Analysis of variance (ANOVA) tests. Data that did not pass the variance test were compared with non-parametric two-tailed Mann-Whitney Rank Sum tests or ANOVA on Ranks tests.

Regression analyses were performed using R v4.0.2 and the “lm” function in the base stats package. Predictor variables considered in regression models were: disease group, age, gender, year of sample, disease duration, disease activity, steroids, biologics, DMARDs and other immunosuppressants, and autoantibody. Age was discretised as <13 or ≥13 based on the distributions of the antibody data by age and the lack of samples from JSLE patients under 13 years of age. All predictor variables were categorical, as described above. Outcome variables modelled were the total, IgG and IgA antibody levels for SARS-CoV-2 and OC43 spike proteins and nucleoproteins. A square-root transformation was applied to the outcome variables in order to better meet assumptions of linearity in regression. Estimates and 95% confidence intervals (CIs) are reported on the square-root scale and do not relate antibody levels on the original scale, however, the antibody levels represent relative fluorescence and do not have meaningful units anyway. The sign of the estimates and the magnitude of estimates within categorical variables are meaningful.

Initially univariable models were fitted for each predictor variable in order to identify which predictors displayed an association with each of the outcome variables. Multiple regression models were subsequently built, starting with a base model comprising disease group, age and gender to ensure age and gender effects were always controlled for. The other predictor variables were added in sequentially, with likelihood ratio tests (LRTs) used to determine which additional variables significantly added to the model. LRTs were performed using the “lrttest” function in the lmtest package v0.9-38. Residual plots were checked to confirm the regression assumptions of linearity and homoscedasticity were met.

## Acknowledgements

We thank L. James and J. Luptak for the SARV-CoV-2 nucleoprotein expression plasmid and M. Pizzato for the SARS-CoV-2 spike expression plasmid. We are grateful for assistance from the Flow Cytometry and Cell Services facilities at the Francis Crick Institute and to Mr Michael Bennet and Mr Simon Caidan for training and support in the high-containment laboratory. We also thank the entire CRICK COVID-19 Consortium. This work was supported by a Centre of Excellence Centre for Adolescent Rheumatology Versus Arthritis grant, 21593, UKRI funding reference MR/R013926/1, the Great Ormond Street Children’s Charity, Cure JM Foundation, Myositis UK, Lupus UK and the NIHR Biomedical Research Centres at GOSH and UCLH. This work was supported by the Francis Crick Institute, which receives its core funding from Cancer Research UK, the UK Medical Research Council, and the Wellcome Trust.

