## Supplementary material for "Favourable antibody responses to human coronaviruses in children and adolescents with autoimmune rheumatic diseases": Figures S1-S3

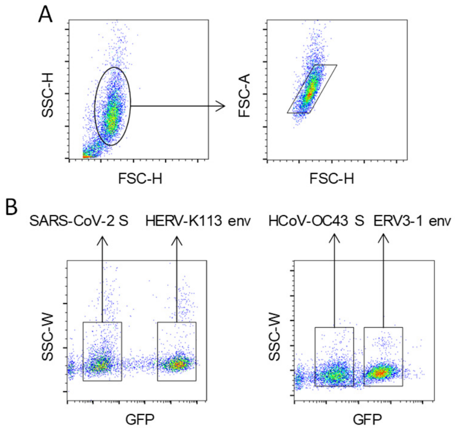


**Figure S1. Gating of cells expressing cell membrane-bound coronaviral spikes or retroviral envelope glycoproteins.** HEK293T cells were transfected with expression plasmids encoding the HCoV-OC43 or SARS-CoV-2 spikes or the ERV3-1 or HERV-K113 envelope glycoproteins and were used for flow cytometric analysis two days later. HEK293T cells expressing HCoV-OC43 spike were mixed with cells expressing ERV3-1 envelope glycoprotein and HEK293T cells expressing SARS-CoV-2 spike were mixed with cells expressing HERV-K113 envelope glycoprotein. (**A**) Gating of HEK293T cells and of single cells in these mixed cell suspensions. (**B**) HEK293T cells expressing the retroviral envelope glycoproteins additionally expressed GFP from the same bicistronic construct and were distinguished from HEK293T cells expressing coronaviral spikes on the basis of GFP expression.

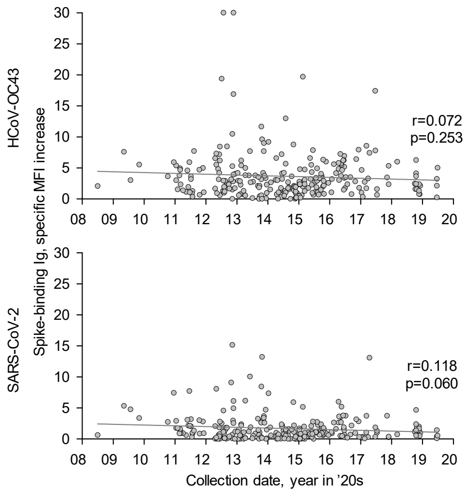


**Figure S2. Effect of sampling date on the titres of antibodies to HCoV-OC43 or SARS-CoV-2 spikes.** The specific increase in MFI of HEK293T cells expressing HCoV-OC43 (top) or SARS-CoV-2 spikes (bottom) is plotted against the date of sample collection. Symbol represent the total IgG, IgM and IgA signal for each spike in individual donors or patients.

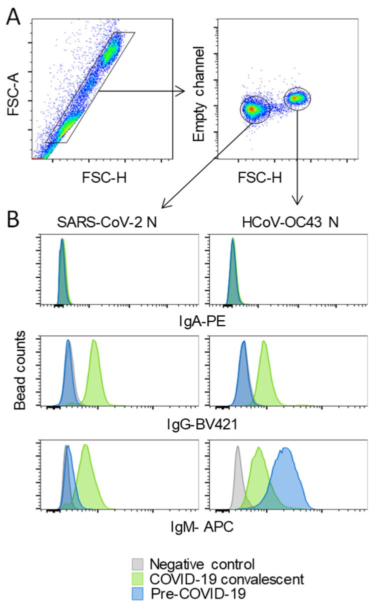


**Figure S3. Detection of antibodies to OC43 or SARS-CoV-2 nucleoproteins by bead-based flow cytometry.** Polymethylmethacrylate beads of different diameter were coated with HCoV-OC43 or SARS-CoV-2 nucleoproteins and were mixed at equal ratios. (**A**) Gating of nucleoprotein-coated beads according to their size. (**B**) Representative staining of beads with a COVID-19 convalescent serum and pre-COVID-19 serum. Unstained beads are also included as a negative control. IgA, IgG and IgM antibodies were detected with the respective secondary antibody.
