## Supplementary material for "Favourable antibody responses to human coronaviruses in children and adolescents with autoimmune rheumatic diseases": Tables S1-S11

**Table S1. Demographic and clinical features of healthy children and adolescents, and patients with JIA, JSLE and JDM**

| Feature | JSLE (n=30) | JIA (n=118) | JDM (n=49) | Control (n=54) |
|---|---|---|---|---|
| Gender, n (%) | | | | |
| Male | 6 (20.0%) | 47 (39.8%) | 18 (36.7%) | 25 (46.3%)[a] |
| Female | 24 (80.0%) | 71 (60.2%) | 31 (63.3%) | 29 (53.7%)[b] |
| Age, median [IQR] | 16 [16-17] | 15 [11-16] | 14 [9-16] | 14 [12-16] |
| Age category, n (%) | | | | |
| Under 13 | 0 (0.0%) | 35 (29.7%) | 24 (49.0%) | 14 (25.9%) |
| Over 13 | 30 (100.0%) | 83 (70.3%) | 25 (51.0%) | 40 (74.1%) |
| Year of sample, n (%) | | | | |
| 2012 | 13 (43.3%) | 18 (15.3%) | 6 (12.2%) | 0 (0.0%) |
| 2013 | 7 (23.3%) | 27 (22.9%) | 4 (8.2%) | 0 (0.0%) |
| 2014 | 5 (16.7%) | 28 (23.7%) | 3 (6.1%) | 4 (7.4%) |
| 2015 | 2 (6.7%) | 15 (12.7%) | 16 (32.7%) | 16 (29.6%) |
| 2016 | 2 (6.7%) | 10 (8.5%) | 4 (8.2%) | 11 (20.4%) |
| Other | 1 (3/3%) | 20 (16.9%) | 16 (32.7%) | 23 (42.6%) |
| Disease duration, n (%) | | | | |
| No disease/diagnosis | 0 (0.0%) | 1 (0.8%) | 0 (0.0%) | 54 (100.0%) |
| Less than 24 months | 12 (40.0%) | 51 (43.2%) | 16 (32.7%) | - |
| Over 24 months | 18 (60.0%) | 66 (55.9%) | 33 (67.3%) | - |
| Disease activity, n (%) | | | | |
| No disease | 7 (23.3%) | 22 (18.6%) | 24 (49.0%) | 54 (100.0%) |
| Mild | 19 (63.3%) | 47 (39.8%) | 18 (36.7%) | - |
| Moderate to severe | 4 (13.3%) | 48 (40.7%) | 7 (14.3%) | - |
| Not assessed | 0 (0.0%) | 1 (0.9%) | 0 (0.0%) | - |
| Autoantibody, n (%) | | | | |
| Negative | 7 (23.3%) | 49 (41.5%) | 11 (22.4%) | - |
| Positive | 23 (76.7%) | 58 (49.2%) | 31 (63.3%) | - |
| Not assessed | 0 (0.0%) | 11 (9.3%) | 7 (14.3%) | 54 (100.0%) |
| Steroid medication, n (%) | | | | |
| No | 12 (40.0%) | 98 (83.1%) | 34 (69.4%) | 54 (100.0%) |
| Yes | 18 (60.0%) | 20 (16.9%) | 15 (30.6%) | - |
| DMARDs and other immunosuppressants, n (%) | | | | |
| No | 2 (6.7%) | 60 (50.8%) | 13 (26.5%) | 54 (100.0%) |
| Yes | 28 (93.3%) | 58 (49.2%) | 36 (73.5%) | - |
| Biologic medication, n (%) | | | | |
| No | 28 (93.3%) | 99 (83.9%) | 46 (93.9%) | 54 (100.0%) |
| Yes | 2 (6.7%) | 19 (16.1%) | 3 (6.1%) | - |

[a]Including n=1 gender dysphoria patient assigned male at birth
[b]Including n=5 gender dysphoria patients assigned female at birth

**Table S2. Univariate models for total, IgG and IgM antibodies to HCoV-OC43 spike protein in healthy children and adolescents, and patients with JIA, JSLE and JDM**

| Predictor[a] | Estimate | 95% CI | p-value | Adjusted $R^2$ |
|---|---|---|---|---|
| ***1. Univariate models for total antibodies to HCoV-OC43 spike protein*** | | | | |
| Disease group | | | | 0.16 |
| Control | - | - | - | |
| JIA | 0.17 | -0.14, 0.43 | 0.21 | |
| JSLE | 1.30 | 0.87, 1.64 | $1.0\times10^{-10}$ | |
| JDM | 0.38 | 0.01, 0.69 | 0.023 | |
| Age | | | | 0 |
| ≤13 | - | - | - | |
| >13 | -0.10 | -0.35, 0.15 | 0.43 | |
| Gender | | | | 0.04 |
| Male | - | - | - | |
| Female | 0.38 | 0.15, 0.61 | 0.0013 | |
| Disease duration | | | | 0.04 |
| No disease/diagnosis | - | - | - | |
| Less than 24 months | 0.53 | 0.22, 0.84 | $8.8\times10^{-4}$ | |
| Over 24 months | 0.32 | 0.03, 0.61 | 0.030 | |
| Disease activity | | | | 0.01 |
| No disease | - | - | - | |
| Mild | 0.32 | 0.05, 0.58 | 0.019 | |
| Moderate to severe | 0.13 | -0.16, 0.43 | 0.36 | |
| Not assessed | -0.28 | -2.07, 1.52 | 0.76 | |
| Year of sample | | | | 0.05 |
| 2012 | - | - | - | |
| 2013 | -0.62 | -1.03, -0.21 | 0.0030 | |
| 2014 | -0.78 | -1.18, -0.38 | $1.4\times10^{-4}$ | |
| 2015 | -0.56 | -0.94, -0.18 | 0.0043 | |
| 2016 | -0.20 | -0.64, 0.25 | 0.39 | |
| Other | -0.47 | -0.83, -0.10 | 0.031 | |
| Steroids | | | | 0.10 |
| No | - | - | - | |
| Yes | 0.71 | 0.45, 0.98 | $3.5\times10^{-7}$ | |
| DMARDs or other immunosuppressants | | | | 0.03 |
| No | - | - | - | |
| Yes | 0.32 | 0.09, 0.55 | 0.0055 | |
| Biologics | | | | 0.01 |
| No | - | - | - | |
| Yes | -0.31 | -0.70, 0.08 | 0.12 | |
| Autoantibody | | | | 0.04 |
| Negative | - | - | - | |
| Positive | 0.39 | 0.16, 0.63 | $9.5\times10^{-4}$ | |
| Not assessed | 0.03 | -0.43, 0.48 | 0.91 | |
| ***2. Univariate models for IgG antibodies to HCoV-OC43 spike protein*** | | | | |
| Disease group | | | | 0.14 |
| Control | - | - | - | |
| JIA | 0.21 | -0.02, 0.45 | 0.074 | |
| JSLE | 1.00 | 0.68, 1.32 | $4.2\times10^{-9}$ | |

| | | | | |
|---|---|---|---|---|
| JDM | 0.53 | 0.25, 0.81 | $2.5\times10^{-4}$ | |
| Age | | | | 0.01 |
| ≤13 | - | - | - | |
| >13 | -0.19 | -0.40, 0.03 | 0.086 | |
| Gender | | | | 0.04 |
| Male | - | - | - | |
| Female | 0.35 | 0.16, 0.55 | $4.7\times10^{-4}$ | |
| Disease duration | | | | 0.05 |
| No disease/diagnosis | - | - | - | |
| Less than 24 months | 0.53 | 0.27, 0.79 | $8.1\times10^{-5}$ | |
| Over 24 months | 0.34 | 0.10, 0.58 | 0.0064 | |
| Disease activity | | | | 0.01 |
| No disease | - | - | - | |
| Mild | 0.28 | 0.06, 0.50 | 0.014 | |
| Moderate to severe | 0.19 | -0.05, 0.44 | 0.13 | |
| Not assessed | -0.09 | -1.61, 1.44 | 0.91 | |
| Year of sample | | | | 0.05 |
| 2012 | - | - | - | |
| 2013 | -0.35 | -0.70, -0.01 | 0.046 | |
| 2014 | -0.54 | -0.88, -0.20 | 0.0019 | |
| 2015 | -0.42 | -0.74, -0.09 | 0.012 | |
| 2016 | 0.06 | -0.32, 0.44 | 0.75 | |
| Other | -0.17 | -0.48, 0.14 | 0.28 | |
| Steroids | | | | 0.08 |
| No | - | - | - | |
| Yes | 0.55 | 0.32, 0.78 | $4.1\times10^{-6}$ | |
| DMARDs or other immunosuppressants | | | | 0.02 |
| No | - | - | - | |
| Yes | 0.23 | 0.04, 0.42 | 0.021 | |
| Biologics | | | | 0 |
| No | - | - | - | |
| Yes | -0.24 | -0.58, 0.09 | 0.15 | |
| Autoantibody | | | | 0.06 |
| Negative | - | - | - | |
| Positive | 0.41 | 0.22, 0.61 | $4.2\times10^{-5}$ | |
| Not assessed | 0.15 | -0.23, 0.54 | 0.44 | |
| ***3. Univariate models for IgM antibodies to HCoV-OC43 spike protein*** | | | | |
| Disease group | | | | 0.11 |
| Control | - | - | - | |
| JIA | -0.06 | -0.21, 0.09 | 0.46 | |
| JSLE | 0.45 | 0.24, 0.66 | $2.9\times10^{-5}$ | |
| JDM | -0.10 | -0.28, 0.08 | 0.26 | |
| Age | | | | 0 |
| ≤13 | - | - | - | |
| >13 | -0.038 | -0.17, 0.10 | 0.57 | |
| Gender | | | | 0 |
| Male | - | - | - | |
| Female | 0.04 | -0.09, 0.16 | 0.57 | |
| Disease duration | | | | 0.01 |
| No disease/diagnosis | - | - | - | |
| Less than 24 months | 0.10 | -0.07, 0.27 | 0.25 | |

| | | | | |
|---|---|---|---|---|
| Over 24 months | -0.04 | -0.20, 0.12 | 0.60 | |
| Disease activity | | | | 0 |
| No disease | - | - | - | |
| Mild | 0.09 | -0.05, 0.23 | 0.22 | |
| Moderate to severe | 0.003 | -0.15, 0.16 | 0.97 | |
| Not assessed | -0.17 | -1.14, 0.80 | 0.73 | |
| Year of sample | | | | 0.04 |
| 2012 | - | - | - | |
| 2013 | -0.33 | -0.55, -0.11 | 0.0031 | |
| 2014 | -0.38 | -0.59, -0.16 | $6.1\times10^{-4}$ | |
| 2015 | -0.26 | -0.46, -0.05 | 0.014 | |
| 2016 | -0.20 | -0.44, 0.03 | 0.093 | |
| Other | -0.31 | -0.51, -0.11 | 0.024 | |
| Steroids | | | | 0.04 |
| No | - | - | - | |
| Yes | 0.25 | 0.10, 0.40 | $9.4\times10^{-4}$ | |
| DMARDs or other immunosuppressants | | | | 0.01 |
| No | - | - | - | |
| Yes | 0.10 | -0.02, 0.22 | 0.11 | |
| Biologics | | | | 0 |
| No | - | - | - | |
| Yes | -0.13 | -0.35, 0.08 | 0.21 | |
| Autoantibody | | | | 0 |
| Negative | - | - | - | |
| Positive | 0.05 | -0.07, 0.18 | 0.40 | |
| Not assessed | -0.08 | -0.33, 0.17 | 0.54 | |

[a]Reference categories for categorical variables indicated by "-"

[b]Sum of IgG, IgM and IgA antibodies

**Table S3. Models for total, IgG and IgM antibodies to HCoV-OC43 spike protein in healthy children and adolescents, and patients with JIA, JSLE and JDM**

| Predictor[a] | Estimate | 95% CI | p-value | Adjusted $R^2$ |
|---|---|---|---|---|
| ***1. Model for total[b] antibodies to HCoV-OC43 spike*** | | | | 0.25 |
| Disease group | | | | |
| Control | - | - | - | |
| JIA | 0.25 | -0.04, 0.55 | 0.094 | |
| JSLE | 1.17 | 0.73, 1.61 | $3.2\times10^{-7}$ | |
| JDM | 0.27 | -0.06, 0.61 | 0.11 | |
| Age | | | | |
| ≤13 | - | - | - | |
| >13 | -0.15 | -0.42, 0.11 | 0.26 | |
| Gender | | | | |
| Male | - | - | - | |
| Female | 0.24 | 0.04, 0.45 | 0.022 | |
| Year of sample | | | | |
| 2012 | - | - | - | |
| 2013 | -0.43 | -0.80, -0.05 | 0.025 | |
| 2014 | -0.43 | -0.80, -0.06 | 0.023 | |
| 2015 | -0.12 | -0.48, 0.26 | 0.56 | |
| 2016 | 0.23 | -0.21, 0.66 | 0.30 | |
| Other | -0.05 | -0.45, 0.35 | 0.81 | |
| Steroids | | | | |
| No | - | - | - | |
| Yes | 0.36 | 0.08, 0.63 | 0.012 | |
| ***2. Model for IgG antibodies to HCoV-OC43 spike*** | | | | 0.24 |
| Disease group | | | | |
| Control | - | - | - | |
| JIA | 0.32 | 0.07, 0.57 | 0.013 | |
| JSLE | 0.99 | 0.61, 1.36 | $4.5\times10^{-7}$ | |
| JDM | 0.49 | 0.20, 0.78 | $8.4\times10^{-4}$ | |
| Age | | | | |
| ≤13 | - | - | - | |
| >13 | -0.11 | -0.33, 0.12 | 0.35 | |
| Gender | | | | |
| Male | - | - | - | |
| Female | 0.23 | 0.05, 0.41 | 0.012 | |
| Year of sample | | | | |
| 2012 | - | - | - | |
| 2013 | -0.20 | -0.52, 0.11 | 0.21 | |
| 2014 | -0.25 | -0.56, 0.06 | 0.12 | |
| 2015 | -0.07 | -0.39, 0.25 | 0.67 | |
| 2016 | 0.42 | 0.06, 0.79 | 0.024 | |
| Other | 0.18 | -0.16, 0.52 | 0.30 | |
| Steroids | | | | |
| No | - | - | - | |
| Yes | 0.27 | 0.04, 0.51 | 0.024 | |

| | | | | |
|---|---|---|---|---|
| ***3. Model for IgM antibodies to HCoV-OC43 spike*** | | | | 0.12 |
| Disease group | | | | |
| Control | - | - | - | |
| JIA | -0.06 | -0.21, 0.09 | 0.42 | |
| JSLE | 0.49 | 0.28, 0.70 | $8.4\times10^{-6}$ | |
| JDM | -0.14 | -0.32, 0.05 | 0.15 | |
| Age | | | | |
| ≤13 | - | - | - | |
| >13 | -0.15 | -0.28, -0.01 | 0.030 | |
| Gender | | | | |
| Male | - | - | - | |
| Female | -0.01 | -0.13, 0.11 | 0.87 | |

[a]Reference categories for categorical variables indicated by "-"

[b]Sum of IgG, IgM and IgA antibodies

**Table S4. Univariate models for total, IgG and IgM antibodies to SARS-CoV-2 spike protein in healthy children and adolescents, and patients with JIA, JSLE and JDM**

| Predictor[a] | Estimate | 95% CI | p-value | Adjusted $R^2$ |
|---|---|---|---|---|
| ***1. Univariate models for total antibodies to SARS-CoV-2 spike protein*** | | | | |
| Disease group | | | | 0.07 |
| Control | - | - | - | |
| JIA | -0.03 | -0.33, 0.14 | 0.79 | |
| JSLE | 0.53 | 0.15, 0.78 | $8.6\times10^{-4}$ | |
| JDM | -0.16 | -0.50, 0.06 | 0.25 | |
| Age | | | | 0.06 |
| ≤13 | - | - | - | |
| >13 | -0.39 | -0.58, -0.20 | $7.2\times10^{-5}$ | |
| Gender | | | | 0.05 |
| Male | - | - | - | |
| Female | 0.33 | 0.15, 0.51 | $3.2\times10^{-4}$ | |
| Disease duration | | | | 0.04 |
| No disease/diagnosis | - | - | - | |
| Less than 24 months | 0.22 | -0.02, 0.46 | 0.075 | |
| Over 24 months | -0.13 | -0.35, 0.10 | 0.27 | |
| Disease activity | | | | 0.03 |
| No disease | - | - | - | |
| Mild | 0.18 | -0.02, 0.38 | 0.081 | |
| Moderate to severe | 0.33 | 0.10, 0.56 | 0.0042 | |
| Not assessed | -0.93 | -2.32, 0.46 | 0.19 | |
| Year of sample | | | | 0.04 |
| 2012 | - | - | - | |
| 2013 | -0.07 | -0.33, 0.31 | -0.97 | |
| 2014 | -0.34 | -0.65, -0.03 | 0.034 | |
| 2015 | -0.17 | -0.47, 0.13 | 0.26 | |
| 2016 | 0.20 | -0.15, 0.55 | 0.26 | |
| Other | 0.13 | -0.16, 0.41 | 0.39 | |
| Steroids | | | | 0 |
| No | - | - | - | |
| Yes | 0.14 | -0.08, 0.36 | 0.20 | |
| DMARDs or other immunosuppressants | | | | 0 |
| No | - | - | - | |
| Yes | -0.06 | -0.23, 0.12 | 0.54 | |
| Biologics | | | | 0.03 |
| No | - | - | - | |
| Yes | -0.45 | -0.75, -0.15 | 0.0037 | |
| Autoantibody | | | | 0.08 |
| Negative | - | - | - | |
| Positive | 0.41 | 0.21, 0.57 | $7.5\times10^{-6}$ | |
| Not assessed | -0.13 | -0.47, 0.22 | 0.47 | |
| ***2. Univariate models for IgG antibodies to SARS-CoV-2 spike protein*** | | | | |
| Disease group | | | | 0.08 |
| Control | - | - | - | |
| JIA | 0.12 | -0.04, 0.35 | 0.12 | |
| JSLE | 0.67 | 0.40, 0.94 | $1.5\times10^{-6}$ | |

| | | | | |
|---|---|---|---|---|
| JDM | 0.12 | -0.12, 0.35 | 0.32 | |
| Age | | | | 0.04 |
| ≤13 | - | - | - | |
| >13 | -0.29 | -0.45, -0.12 | $9.9\times10^{-4}$ | |
| Gender | | | | 0.06 |
| Male | - | - | - | |
| Female | 0.33 | 0.17, 0.49 | $4.5\times10^{-5}$ | |
| Disease duration | | | | 0.05 |
| No disease/diagnosis | - | - | - | |
| Less than 24 months | 0.38 | 0.16, 0.59 | $5.4\times10^{-4}$ | |
| Over 24 months | 0.11 | -0.09, 0.31 | 0.28 | |
| Disease activity | | | | 0.05 |
| No disease | - | - | - | |
| Mild | 0.21 | 0.04, 0.39 | 0.017 | |
| Moderate to severe | 0.35 | 0.16, 0.55 | $4.6\times10^{-4}$ | |
| Not assessed | -0.66 | -1.87, 0.55 | 0.29 | |
| Year of sample | | | | 0.04 |
| 2012 | - | - | - | |
| 2013 | -0.02 | -0.30, 0.26 | 0.90 | |
| 2014 | -0.32 | -0.60, -0.05 | 0.021 | |
| 2015 | -0.25 | -0.52, 0.01 | 0.059 | |
| 2016 | 0.11 | -0.20, 0.41 | 0.48 | |
| Other | 0.04 | -0.22, 0.29 | 0.77 | |
| Steroids | | | | 0.01 |
| No | - | - | - | |
| Yes | 0.17 | -0.02, 0.36 | 0.084 | |
| DMARDs or other immunosuppressants | | | | 0 |
| No | - | - | - | |
| Yes | 0.04 | -0.12, 0.19 | 0.66 | |
| Biologics | | | | 0.02 |
| No | - | - | - | |
| Yes | -0.33 | -0.59, -0.06 | 0.017 | |
| Autoantibody | | | | 0.13 |
| Negative | - | - | - | |
| Positive | 0.47 | 0.32, 0.62 | $4.5\times10^{-9}$ | |
| Not assessed | 0.02 | -0.28, 0.32 | 0.90 | |
| ***3. Univariate models for IgM antibodies to SARS-CoV-2 spike protein*** | | | | |
| Disease group | | | | 0.13 |
| Control | - | - | - | |
| JIA | -0.23 | -0.36, -0.11 | $3.6\times10^{-4}$ | |
| JSLE | -0.04 | -0.22, 0.13 | 0.62 | |
| JDM | -0.45 | -0.60, -0.30 | $1.5\times10^{-8}$ | |
| Age | | | | 0.06 |
| ≤13 | - | - | - | |
| >13 | -0.24 | -0.35, -0.13 | $2.9\times10^{-5}$ | |
| Gender | | | | 0.01 |
| Male | - | - | - | |
| Female | 0.08 | -0.03, 0.19 | 0.14 | |
| Disease duration | | | | 0.09 |
| No disease/diagnosis | - | - | - | |
| Less than 24 months | -0.16 | -0.29, -0.02 | 0.024 | |

| | | | | |
|---|---|---|---|---|
| Over 24 months | -0.33 | -0.46, -0.20 | $6.1\times10^{-7}$ | |
| Disease activity | | | | 0 |
| No disease | - | - | - | |
| Mild | 0.002 | -0.12, 0.12 | 0.97 | |
| Moderate to severe | 0.06 | -0.07, 0.19 | 0.39 | |
| Not assessed | -0.51 | -1.33, 0.32 | 0.23 | |
| Year of sample | | | | 0.03 |
| 2012 | - | - | - | |
| 2013 | 0.04 | -0.15, 0.23 | 0.66 | |
| 2014 | -0.09 | -0.27, 0.10 | 0.36 | |
| 2015 | 0.02 | -0.15, 0.20 | 0.80 | |
| 2016 | 0.19 | -0.02, 0.39 | 0.070 | |
| Other | 0.17 | 0.00, 0.39 | 0.045 | |
| Steroids | | | | 0 |
| No | - | - | - | |
| Yes | -0.05 | -0.18, 0.07 | 0.41 | |
| DMARDs or other immunosuppressants | | | | 0.05 |
| No | - | - | - | |
| Yes | -0.19 | -0.29, -0.09 | $3.4\times10^{-4}$ | |
| Biologics | | | | 0.04 |
| No | - | - | - | |
| Yes | -0.30 | -0.48, -0.13 | $7.5\times10^{-4}$ | |
| Autoantibody | | | | 0.02 |
| Negative | - | - | - | |
| Positive | 0.04 | -0.07, 0.14 | 0.52 | |
| Not assessed | -0.25 | -0.46, -0.04 | 0.022 | |

[a]Reference categories for categorical variables indicated by "-"

[b]Sum of IgG, IgM and IgA antibodies

**Table S5. Models for total, IgG and IgM antibodies to SARS-CoV-2 spike protein in healthy children and adolescents, and patients with JIA, JSLE and JDM**

| Predictor[a] | Estimate | 95% CI | p-value | Adjusted $R^2$ |
|---|---|---|---|---|
| ***1. Model for total[b] antibodies to SARS-CoV-2 spike*** | | | | 0.26 |
| Disease group | | | | |
| Control | - | - | - | |
| JIA | -0.23 | -0.47, -0.002 | 0.048 | |
| JSLE | 0.32 | 0.002, 0.64 | 0.049 | |
| JDM | -0.54 | -0.82, -0.26 | $1.9\times10^{-4}$ | |
| Age | | | | |
| ≤13 | - | - | - | |
| >13 | -0.48 | -0.66, -0.30 | $3.0\times10^{-7}$ | |
| Gender | | | | |
| Male | - | - | - | |
| Female | 0.18 | 0.01, 0.34 | 0.036 | |
| Biologics | | | | |
| No | - | - | - | |
| Yes | -0.25 | -0.54, 0.03 | 0.083 | |
| Autoantibody | | | | |
| Negative | - | - | - | |
| Positive | 0.40 | 0.20, 0.59 | $1.1\times10^{-4}$ | |
| Not assessed | 0.15 | -0.19, 0.49 | 0.39 | |
| ***2. Model for IgG antibodies to SARS-CoV-2 spike*** | | | | 0.27 |
| Disease group | | | | |
| Control | - | - | - | |
| JIA | 0.02 | -0.20, 0.24 | 0.85 | |
| JSLE | 0.52 | 0.21, 0.82 | 0.0011 | |
| JDM | -0.17 | -0.42, 0.08 | 0.18 | |
| Age | | | | |
| ≤13 | - | - | - | |
| >13 | -0.27 | -0.45, -0.09 | 0.010 | |
| Gender | | | | |
| Male | - | - | - | |
| Female | 0.16 | 0.01, 0.30 | 0.023 | |
| Year of sample | | | | |
| 2012 | - | - | - | |
| 2013 | -0.03 | -0.28, 0.22 | 0.82 | |
| 2014 | -0.13 | -0.38, 0.11 | 0.29 | |
| 2015 | -0.03 | -0.28, 0.23 | 0.84 | |
| 2016 | 0.29 | -0.001, 0.58 | 0.051 | |
| Other | 0.13 | -0.14, 0.40 | 0.34 | |
| Autoantibody | | | | |
| Negative | - | - | - | |
| Positive | 0.38 | 0.20, 0.56 | $3.1\times10^{-5}$ | |
| Not assessed | 0.15 | -0.15, 0.44 | 0.33 | |
| ***3. Model for IgM antibodies to SARS-CoV-2 spike*** | | | | 0.30 |
| Disease group | | | | |
| Control | - | - | - | |

| | | | |
|---|---|---|---|
| JIA | -0.30 | -0.43, -0.17 | $1.0\times10^{-5}$ |
| JSLE | -0.08 | -0.26, 0.10 | 0.37 |
| JDM | -0.62 | -0.78, -0.46 | $7.3\times10^{-13}$ |
| Age | | | |
| ≤13 | - | - | - |
| >13 | -0.31 | -0.42, -0.21 | $5.9\times10^{-9}$ |
| Gender | | | |
| Male | - | - | - |
| Female | 0.03 | -0.06, 0.13 | 0.52 |
| Biologics | | | |
| No | - | - | - |
| Yes | -0.15 | -0.31, 0.01 | 0.071 |
| Autoantibody | | | |
| Negative | - | - | - |
| Positive | 0.16 | 0.05, 0.27 | 0.0062 |
| Not assessed | 0.01 | -0.19, 0.20 | 0.95 |

[a]Reference categories for categorical variables indicated by "-"

[b]Sum of IgG, IgM and IgA antibodies

**Table S6. Univariate models for total, IgG and IgM antibodies to HCoV-OC43 nucleoprotein in healthy children and adolescents, and patients with JIA, JSLE and JDM**

| Predictor[a] | Estimate | 95% CI | p-value | Adjusted $R^2$ |
|---|---|---|---|---|
| ***1. Univariate models for total antibodies to HCoV-OC43 nucleoprotein*** | | | | |
| Disease group | | | | 0.30 |
| Control | - | - | - | |
| JIA | -0.30 | -0.59, -0.02 | 0.036 | |
| JSLE | 0.48 | 0.09, 0.88 | 0.016 | |
| JDM | -1.40 | -1.76 -1.07 | $1.8\times10^{-14}$ | |
| Age | | | | 0 |
| ≤13 | - | - | - | |
| >13 | 0.19 | -0.10, 0.47 | 0.20 | |
| Gender | | | | 0.01 |
| Male | - | - | - | |
| Female | -0.23 | -0.49, 0.04 | 0.098 | |
| Disease duration | | | | 0.04 |
| No disease/diagnosis | - | - | - | |
| Less than 24 months | -0.29 | -0.65, 0.06 | 0.11 | |
| Over 24 months | -0.57 | -0.90, -0.24 | $7.6\times10^{-4}$ | |
| Disease activity | | | | 0 |
| No disease | - | - | - | |
| Mild | 0.05 | -0.25, 0.36 | 0.72 | |
| Moderate to severe | -0.06 | -0.40, 0.28 | 0.72 | |
| Not assessed | 0.69 | -1.39, 2.77 | 0.51 | |
| Year of sample | | | | 0 |
| 2012 | - | - | - | |
| 2013 | -0.16 | -0.63, 0.32 | 0.52 | |
| 2014 | -0.03 | -0.50, 0.45 | 0.91 | |
| 2015 | -0.06 | -0.51, 0.40 | 0.81 | |
| 2016 | 0.25 | -0.28, 0.77 | 0.35 | |
| Other | -0.02 | -0.45, 0.41 | 0.93 | |
| Steroids | | | | 0 |
| No | - | - | - | |
| Yes | 0.22 | -0.10, 0.53 | 0.18 | |
| DMARDs or other immunosuppressants | | | | 0 |
| No | - | - | - | |
| Yes | -0.00078 | -0.26, 0.26 | 1.0 | |
| Biologics | | | | 0 |
| No | - | - | - | |
| Yes | 0.19 | -0.25, 0.64 | 0.39 | |
| Autoantibody | | | | 0.04 |
| Negative | - | - | - | |
| Positive | -0.45 | -0.71, -0.18 | 0.0011 | |
| Not assessed | -0.33 | -0.84, 0.18 | 0.21 | |
| ***2. Univariate models for IgG antibodies to HCoV-OC43 nucleoprotein*** | | | | |
| Disease group | | | | 0.03 |
| Control | - | - | - | |
| JIA | -0.07 | -0.37, 0.22 | 0.63 | |
| JSLE | 0.54 | 0.13, 0.95 | 0.010 | |

| | | | | |
|---|---|---|---|---|
| JDM | 0.02 | -0.39, 0.43 | 0.92 | |
| Age | | | | 0 |
| ≤13 | - | - | - | |
| >13 | 0.12 | -0.14, 0.38 | 0.36 | |
| Gender | | | | 0.01 |
| Male | - | - | - | |
| Female | 0.19 | -0.06, 0.43 | 0.13 | |
| Disease duration | | | | 0 |
| No disease/diagnosis | - | - | - | |
| Less than 24 months | 0.03 | -0.30, 0.35 | 0.87 | |
| Over 24 months | 0.04 | -0.27, 0.33 | 0.80 | |
| Disease activity | | | | 0 |
| No disease | - | - | - | |
| Mild | 0.03 | -0.25, 0.31 | 0.83 | |
| Moderate to severe | -0.06 | -0.37, 0.25 | 0.70 | |
| Not assessed | -0.39 | -2.24, 1.46 | 0.68 | |
| Year of sample | | | | 0 |
| 2012 | - | - | - | |
| 2013 | -0.19 | -0.65, 0.26 | -0.40 | |
| 2014 | -0.26 | -0.71, 0.18 | 0.25 | |
| 2015 | -0.13 | -0.56, 0.31 | 0.57 | |
| 2016 | 0.17 | -0.31, 0.65 | 0.48 | |
| Other | -0.12 | -0.53, 0.28 | 0.55 | |
| Steroids | | | | 0.03 |
| No | - | - | - | |
| Yes | 0.39 | 0.10, 0.68 | 0.0085 | |
| DMARDs or other immunosuppressants | | | | 0 |
| No | - | - | - | |
| Yes | 0.14 | -0.10, 0.38 | 0.24 | |
| Biologics | | | | 0 |
| No | - | - | - | |
| Yes | -0.15 | -0.55, 0.26 | 0.48 | |
| Autoantibody | | | | 0 |
| Negative | - | - | - | |
| Positive | 0.08 | -0.17, 0.34 | 0.52 | |
| Not assessed | -0.15 | -0.61, 0.32 | 0.54 | |
| ***3. Univariate models for IgM antibodies to HCoV-OC43 nucleoprotein*** | | | | |
| Disease group | | | | 0.13 |
| Control | - | - | - | |
| JIA | -0.27 | -0.48, -0.06 | 0.012 | |
| JSLE | 0.26 | -0.03, 0.55 | 0.083 | |
| JDM | -0.70 | -1.00, -0.41 | $4.1\times10^{-6}$ | |
| Age | | | | 0.08 |
| ≤13 | - | - | - | |
| >13 | 0.42 | 0.23, 0.61 | $1.4\times10^{-5}$ | |
| Gender | | | | 0.02 |
| Male | - | - | - | |
| Female | -0.21 | -0.39, -0.03 | 0.025 | |
| Disease duration | | | | 0.01 |
| No disease/diagnosis | - | - | - | |
| Less than 24 months | -0.24 | -0.48, 0.00 | 0.048 | |

| | | | | |
|---|---|---|---|---|
| Over 24 months | -0.22 | -0.45, 0.01 | 0.064 | |
| Disease activity | | | | 0.01 |
| No disease | - | - | - | |
| Mild | -0.04 | -0.26, 0.16 | 0.74 | |
| Moderate to severe | -0.22 | -0.47, 0.00 | 0.057 | |
| Not assessed | 0.67 | -0.73, 2.03 | 0.34 | |
| Year of sample | | | | 0.03 |
| 2012 | - | - | - | |
| 2013 | -0.20 | -0.53, 0.14 | 0.25 | |
| 2014 | -0.13 | -0.46, 0.19 | 0.42 | |
| 2015 | -0.05 | -0.37, 0.27 | 0.78 | |
| 2016 | -0.34 | -0.69, 0.02 | 0.065 | |
| Other | -0.42 | -0.72, -0.12 | 0.0064 | |
| Steroids | | | | 0 |
| No | - | - | - | |
| Yes | -0.07 | -0.29, 0.15 | 0.55 | |
| DMARDs or other immunosuppressants | | | | 0 |
| No | - | - | - | |
| Yes | 0.06 | -0.12, 0.24 | 0.53 | |
| Biologics | | | | 0 |
| No | - | - | - | |
| Yes | 0.18 | -0.12, 0.48 | 0.23 | |
| Autoantibody | | | | 0.01 |
| Negative | - | - | - | |
| Positive | -0.18 | -0.37, 0.01 | 0.062 | |
| Not assessed | -0.28 | -0.62, 0.07 | 0.11 | |

[a]Reference categories for categorical variables indicated by "-"
[b]Sum of IgG, IgM and IgA antibodies

**Table S7. Models for total, IgG and IgM antibodies to HCoV-OC43 nucleoprotein in healthy children and adolescents, and patients with JIA, JSLE and JDM**

| Predictor[a] | Estimate | 95% CI | p-value | Adjusted $R^2$ |
|---|---|---|---|---|
| ***1. Model for total antibodies to HCoV-OC43 nucleoprotein*** | | | | 0.32 |
| Disease group | | | | |
| Control | - | - | - | |
| JIA | -0.44 | -0.75, -0.13 | 0.0053 | |
| JSLE | 0.34 | -0.13, 0.80 | 0.15 | |
| JDM | -1.67 | -2.07, -1.27 | $1.5\times10^{-14}$ | |
| Age | | | | |
| ≤13 | - | - | - | |
| >13 | -0.25 | -0.50, 0.003 | 0.053 | |
| Gender | | | | |
| Male | - | - | - | |
| Female | -0.26 | -0.49, -0.04 | 0.022 | |
| DMARDs or other immunosuppressants | | | | |
| No | - | - | - | |
| Yes | 0.30 | 0.03, 0.57 | 0.030 | |
| ***2. Model for IgG antibodies to HCoV-OC43 nucleoprotein*** | | | | 0.03 |
| Disease group | | | | |
| Control | - | - | - | |
| JIA | -0.08 | -0.37, 0.22 | 0.60 | |
| JSLE | 0.49 | 0.06, 0.91 | 0.024 | |
| JDM | 0.05 | -0.39, 0.22 | 0.81 | |
| Age | | | | |
| ≤13 | - | - | - | |
| >13 | 0.05 | -0.24, 0.34 | 0.71 | |
| Gender | | | | |
| Male | - | - | - | |
| Female | 0.14 | -0.11, 0.39 | 0.26 | |
| ***3. Model for IgM antibodies to HCoV-OC43 nucleoprotein*** | | | | 0.18 |
| Disease group | | | | |
| Control | - | - | - | |
| JIA | -0.24 | -0.45, -0.04 | 0.019 | |
| JSLE | 0.28 | -0.02, 0.57 | 0.065 | |
| JDM | -0.60 | -0.91, -0.29 | $1.6\times10^{-4}$ | |
| Age | | | | |
| ≤13 | - | - | - | |
| >13 | 0.19 | -0.01, 0.39 | 0.061 | |
| Gender | | | | |
| Male | - | - | - | |
| Female | -0.26 | -0.43, -0.09 | 0.0032 | |

[a]Reference categories for categorical variables indicated by "-"

[b]Sum of IgG, IgM and IgA antibodies

**Table S8. Univariate models for total, IgG and IgM antibodies to SARS-CoV-2 nucleoprotein in healthy children and adolescents, and patients with JIA, JSLE and JDM**

| Predictor[a] | Estimate | 95% CI | p-value | Adjusted $R^2$ |
|---|---|---|---|---|
| ***1. Univariate models for total antibodies to SARS-CoV-2 nucleoprotein*** | | | | |
| Disease group | | | | 0.25 |
| Control | - | - | - | |
| JIA | -0.35 | -0.52, -0.18 | $8.2\times10^{-5}$ | |
| JSLE | 0.28 | 0.04, 0.52 | $2.3\times10^{-2}$ | |
| JDM | -0.75 | -0.96, -0.54 | $9.6\times10^{-12}$ | |
| Age | | | | 0.04 |
| ≤13 | - | - | - | |
| >13 | 0.28 | 0.11, 0.45 | $9.9\times10^{-4}$ | |
| Gender | | | | 0 |
| Male | - | - | - | |
| Female | 0.005 | -0.15, 0.16 | 0.95 | |
| Disease duration | | | | 0.06 |
| No disease/diagnosis | - | - | - | |
| Less than 24 months | -0.28 | -0.48, -0.07 | 0.0090 | |
| Over 24 months | -0.40 | -0.59, -0.21 | $6.3\times10^{-5}$ | |
| Disease activity | | | | 0 |
| No disease | - | - | - | |
| Mild | -0.05 | -0.22, 0.13 | 0.61 | |
| Moderate to severe | -0.01 | -0.21, 0.18 | 0.89 | |
| Not assessed | 0.27 | -0.96, 1.49 | 0.67 | |
| Year of sample | | | | 0 |
| 2012 | - | - | - | |
| 2013 | -0.20 | -0.48, 0.08 | 0.16 | |
| 2014 | -0.03 | -0.31, 0.25 | 0.82 | |
| 2015 | 0.02 | -0.25, 0.28 | 0.89 | |
| 2016 | 0.04 | -0.27, 0.35 | 0.80 | |
| Other | -0.17 | -0.43, 0.08 | 0.18 | |
| Steroids | | | | 0.01 |
| No | - | - | - | |
| Yes | 0.15 | -0.04, 0.34 | 0.11 | |
| DMARDs or other immunosuppressants | | | | 0 |
| No | - | - | - | |
| Yes | -0.11 | -0.27, 0.04 | 0.14 | |
| Biologics | | | | 0 |
| No | - | - | - | |
| Yes | -0.004 | -0.27, 0.26 | 0.97 | |
| Autoantibody | | | | 0.01 |
| Negative | - | - | - | |
| Positive | -0.17 | -0.32, -0.01 | 0.041 | |
| Not assessed | -0.12 | -0.43, 0.18 | 0.43 | |
| ***2. Univariate models for IgG antibodies to SARS-CoV-2 nucleoprotein*** | | | | |
| Disease group | | | | 0.21 |
| Control | - | - | - | |
| JIA | -0.26 | -0.44, -0.20 | $1.2\times10^{-5}$ | |
| JSLE | 0.28 | 0.06, 0.38 | $6.9\times10^{-4}$ | |

| | | | | |
|---|---|---|---|---|
| JDM | -0.20 | -0.43, -0.10 | 0.0015 | |
| Age | | | | 0.07 |
| ≤13 | - | - | - | |
| >13 | 0.24 | 0.14, 0.35 | $1.5\times10^{-5}$ | |
| Gender | | | | 0.02 |
| Male | - | - | - | |
| Female | 0.13 | 0.02, 0.24 | 0.016 | |
| Disease duration | | | | 0.02 |
| No disease/diagnosis | - | - | - | |
| Less than 24 months | -0.18 | -0.32, -0.04 | 0.011 | |
| Over 24 months | -0.14 | -0.27, 0.00 | 0.043 | |
| Disease activity | | | | 0 |
| No disease | - | - | - | |
| Mild | -0.07 | -0.19, 0.05 | 0.25 | |
| Moderate to severe | -0.08 | -0.21, 0.05 | 0.24 | |
| Not assessed | 0.028 | -0.78, 0.83 | 0.95 | |
| Year of sample | | | | 0.01 |
| 2012 | - | - | - | |
| 2013 | -0.20 | -0.39, 0.00 | 0.050 | |
| 2014 | -0.05 | -0.24, 0.15 | 0.63 | |
| 2015 | -0.01 | -0.20, 0.18 | 0.92 | |
| 2016 | -0.10 | -0.30, 0.11 | 0.37 | |
| Other | -0.16 | -0.33, 0.02 | 0.076 | |
| Steroids | | | | 0 |
| No | - | - | - | |
| Yes | 0.11 | -0.02, 0.23 | 0.10 | |
| DMARDs or other immunosuppressants | | | | 0 |
| No | - | - | - | |
| Yes | -0.03 | -0.13, 0.08 | 0.61 | |
| Biologics | | | | 0 |
| No | - | - | - | |
| Yes | -0.08 | -0.25, 0.10 | 0.37 | |
| Autoantibody | | | | 0 |
| Negative | - | - | - | |
| Positive | 0.019 | -0.09, 0.13 | 0.73 | |
| Not assessed | -0.08 | -0.28, 0.12 | 0.43 | |
| ***3. Univariate models for IgM antibodies to SARS-CoV-2 nucleoprotein*** | | | | |
| Disease group | | | | 0.10 |
| Control | - | - | - | |
| JIA | -0.18 | -0.27, 0.04 | 0.0031 | |
| JSLE | 0.10 | 0.18, 0.60 | 0.24 | |
| JDM | -0.31 | -0.34, 0.03 | $2.3\times10^{-4}$ | |
| Age | | | | 0.08 |
| ≤13 | - | - | - | |
| >13 | 0.23 | 0.13, 0.33 | $1.0\times10^{-5}$ | |
| Gender | | | | 0 |
| Male | - | - | - | |
| Female | -0.02 | -0.12, 0.08 | 0.72 | |
| Disease duration | | | | 0.02 |
| No disease/diagnosis | - | - | - | |
| Less than 24 months | -0.14 | -0.27, -0.01 | 0.038 | |

| | | | | |
|---|---|---|---|---|
| Over 24 months | -0.17 | -0.29, -0.05 | 0.0073 | |
| Disease activity | | | | 0 |
| No disease | - | - | - | |
| Mild | -0.05 | -0.08, 0.21 | 0.06 | |
| Moderate to severe | -0.03 | -0.18, 0.14 | 0.62 | |
| Not assessed | 0.19 | -1.16, 0.77 | 0.63 | |
| Year of sample | | | | 0.05 |
| 2012 | - | - | - | |
| 2013 | -0.12 | -0.30, 0.06 | 0.20 | |
| 2014 | -0.10 | -0.28, 0.07 | 0.26 | |
| 2015 | 0.02 | -0.15, 0.20 | 0.79 | |
| 2016 | -0.06 | -0.25, 0.13 | 0.55 | |
| Other | -0.24 | -0.40, -0.08 | 0.0032 | |
| Steroids | | | | 0 |
| No | - | - | - | |
| Yes | 0.04 | -0.08, 0.16 | 0.47 | |
| DMARDs or other immunosuppressants | | | | 0 |
| No | - | - | - | |
| Yes | -0.06 | -0.16, 0.03 | 0.20 | |
| Biologics | | | | 0 |
| No | - | - | - | |
| Yes | -0.02 | -0.19, 0.14 | 0.80 | |
| Autoantibody | | | | 0 |
| Negative | - | - | - | |
| Positive | -0.04 | -0.15, 0.06 | 0.42 | |
| Not assessed | -0.11 | -0.30, 0.08 | 0.26 | |

[a]Reference categories for categorical variables indicated by “-“

[b]Sum of IgG, IgM and IgA antibodies

**Table S9. Models for total, IgG and IgM antibodies to SARS-CoV-2 nucleoprotein in healthy children and adolescents, and patients with JIA, JSLE and JDM**

| Predictor[a] | Estimate | 95% CI | p-value | Adjusted $R^2$ |
|---|---|---|---|---|
| ***1. Model for total antibodies to SARS-CoV-2 nucleoprotein*** | | | | 0.25 |
| Disease group | | | | |
| Control | - | - | - | |
| JIA | -0.34 | -0.52, -0.17 | $1.1\times10^{-4}$ | |
| JSLE | 0.26 | 0.01, 0.50 | 0.039 | |
| JDM | -0.73 | -0.94, -0.52 | $7.3\times10^{-11}$ | |
| Age | | | | |
| ≤13 | - | - | - | |
| >13 | 0.09 | -0.06, 0.25 | 0.22 | |
| Gender | | | | |
| Male | - | - | - | |
| Female | -0.02 | -0.16, 0.12 | 0.79 | |
| ***2. Model for IgG antibodies to SARS-CoV-2 nucleoprotein*** | | | | 0.27 |
| Disease group | | | | |
| Control | - | - | - | |
| JIA | -0.19 | -0.32, -0.07 | 0.0031 | |
| JSLE | 0.33 | 0.14, 0.52 | $7.9\times10^{-4}$ | |
| JDM | 0.05 | -0.15, 0.25 | 0.64 | |
| Age | | | | |
| ≤13 | - | - | - | |
| >13 | 0.23 | 0.12, 0.35 | $7.0\times10^{-5}$ | |
| Gender | | | | |
| Male | - | - | - | |
| Female | 0.11 | 0.02, 0.21 | 0.019 | |
| DMARDs or other immunosuppressants | | | | |
| No | - | - | - | |
| Yes | -0.15 | -0.26, -0.03 | 0.016 | |
| ***3. Model for IgM antibodies to SARS-CoV-2 nucleoprotein*** | | | | 0.13 |
| Disease group | | | | |
| Control | - | - | - | |
| JIA | -0.18 | -0.29, -0.06 | 0.0029 | |
| JSLE | 0.06 | -0.10, 0.23 | 0.45 | |
| JDM | -0.22 | -0.39, -0.04 | 0.014 | |
| Age | | | | |
| ≤13 | - | - | - | |
| >13 | 0.16 | 0.05, 0.27 | 0.0047 | |
| Gender | | | | |
| Male | - | - | - | |
| Female | -0.03 | -0.12, 0.07 | 0.56 | |

[a]Reference categories for categorical variables indicated by “-“
[b]Sum of IgG, IgM and IgA antibodies

**Table S10. Univariate models for ratio of antibodies to spike and nucleoprotein for HCoV-OC43 and SARS-CoV-2 in healthy children and adolescents, and patients with JIA, JSLE and JDM**

| Predictor[a] | Estimate | 95% CI | p-value | Adjusted $R^2$ |
|---|---|---|---|---|
| ***1. Univariate models for ratio of antibodies to spike and nucleoprotein for HCoV-OC43*** | | | | |
| Disease group | | | | 0.04 |
| Control | - | - | - | |
| JIA | 0.39 | 0.04, 0.36 | 0.071 | |
| JSLE | 0.17 | 0.26, 0.70 | 0.56 | |
| JDM | 1.00 | 0.47, 0.92 | 0.0012 | |
| Age | | | | 0.17 |
| ≤13 | - | - | - | |
| >13 | -1.20 | -1.50, -0.82 | $1.7 \times 10^{-10}$ | |
| Gender | | | | 0.03 |
| Male | - | - | - | |
| Female | 0.48 | 0.13, 0.83 | 0.0077 | |
| Disease duration | | | | 0.03 |
| No disease/diagnosis | - | - | - | |
| Less than 24 months | 0.66 | 0.21, 1.11 | 0.0046 | |
| Over 24 months | 0.26 | -0.18, 0.69 | 0.25 | |
| Disease activity | | | | 0.02 |
| No disease | - | - | - | |
| Mild | 0.45 | 0.05, 0.85 | 0.027 | |
| Moderate to severe | 0.51 | 0.08, 0.94 | 0.022 | |
| Not assessed | -1.10 | -3.62, 1.50 | 0.41 | |
| Year of sample | | | | 0.06 |
| 2012 | - | - | - | |
| 2013 | 0.43 | -0.20, 1.06 | 0.18 | |
| 2014 | -0.32 | -0.94, 0.30 | 0.31 | |
| 2015 | -0.21 | -0.81, 0.39 | 0.49 | |
| 2016 | 0.37 | -0.30, 1.04 | 0.28 | |
| Other | 0.61 | 0.05, 1.18 | 0.034 | |
| Steroids | | | | 0 |
| No | - | - | - | |
| Yes | -0.12 | -0.55, 0.30 | 0.57 | |
| DMARDs or other immunosuppressants | | | | 0 |
| No | - | - | - | |
| Yes | 0.20 | -0.14, 0.55 | 0.24 | |
| Biologics | | | | 0.01 |
| No | - | - | - | |
| Yes | -0.53 | -0.14, 0.55 | 0.069 | |
| Autoantibody | | | | 0.06 |
| Negative | - | - | - | |
| Positive | 0.70 | 0.35, 1.05 | $1.1 \times 10^{-4}$ | |
| Not assessed | 0.33 | -0.32, 0.98 | 0.32 | |
| ***2. Univariate models for ratio of antibodies to spike and nucleoprotein for SARS-CoV-2*** | | | | |
| Disease group | | | | 0.16 |
| Control | - | - | - | |
| JIA | 0.19 | 0.04, 0.35 | 0.013 | |
| JSLE | 0.47 | 0.26, 0.68 | $1.9 \times 10^{-5}$ | |

| | | | | |
|---|---|---|---|---|
| JDM | 0.68 | 0.47, 0.90 | $2.5\times10^{-9}$ | |
| Age | | | | 0.05 |
| ≤13 | - | - | - | |
| >13 | -0.26 | -0.40, -0.112 | $4.4\times10^{-4}$ | |
| Gender | | | | 0.04 |
| Male | - | - | - | |
| Female | 0.22 | 0.09, 0.36 | 0.0015 | |
| Disease duration | | | | 0.07 |
| No disease/diagnosis | - | - | - | |
| Less than 24 months | 0.38 | 0.21, 0.56 | $1.8\times10^{-5}$ | |
| Over 24 months | 0.28 | 0.11, 0.44 | 0.0011 | |
| Disease activity | | | | 0.03 |
| No disease | - | - | - | |
| Mild | 0.25 | 0.09, 0.40 | 0.0017 | |
| Moderate to severe | 0.14 | -0.03, 0.31 | 0.10 | |
| Not assessed | -0.26 | -1.26, 0.75 | 0.62 | |
| Year of sample | | | | 0.03 |
| 2012 | - | - | - | |
| 2013 | -0.16 | -0.41, 0.09 | 0.21 | |
| 2014 | -0.37 | -0.61, -0.13 | 0.0032 | |
| 2015 | -0.21 | -0.45, 0.02 | 0.077 | |
| 2016 | -0.03 | -0.29, 0.23 | 0.83 | |
| Other | -0.13 | -0.35, 0.09 | 0.24 | |
| Steroids | | | | 0.09 |
| No | - | - | - | |
| Yes | 0.39 | 0.23, 0.55 | $2.1\times10^{-6}$ | |
| DMARDs or other immunosuppressants | | | | 0.02 |
| No | - | - | - | |
| Yes | 0.15 | 0.01, 0.28 | 0.030 | |
| Biologics | | | | 0 |
| No | - | - | - | |
| Yes | -0.15 | -0.38, 0.08 | 0.19 | |
| Autoantibody | | | | 0.08 |
| Negative | - | - | - | |
| Positive | 0.32 | 0.19, 0.46 | $4.7\times10^{-6}$ | |
| Not assessed | 0.17 | -0.09, 0.42 | 0.19 | |

[a]Reference categories for categorical variables indicated by "-"

**Table S11. Models for ratio of antibodies to spike and nucleoprotein for HCoV-OC43 and SARS-CoV-2 in healthy children and adolescents, and patients with JIA, JSLE and JDM**

| Predictor[a] | Estimate | 95% CI | p-value | Adjusted $R^2$ |
|---|---|---|---|---|
| ***1. Model for ratio of antibodies to spike and antibodies to nucleoprotein for HCoV-OC43*** | | | | 0.20 |
| Disease group | | | | |
| Control | - | - | - | |
| JIA | 0.09 | -0.36, 0.53 | 0.71 | |
| JSLE | 0.02 | -0.60, 0.64 | 0.96 | |
| JDM | 0.15 | -0.48, 0.79 | 0.64 | |
| Age | | | | |
| ≤13 | - | - | - | |
| >13 | -1.05 | -1.43, -0.67 | $1.5\times10^{-7}$ | |
| Gender | | | | |
| Male | - | - | - | |
| Female | 0.32 | -0.01, 0.66 | 0.060 | |
| Autoantibody | | | | |
| Negative | - | - | - | |
| Positive | 0.46 | 0.06, 0.86 | 0.026 | |
| Not assessed | 0.22 | -0.43, 0.86 | 0.51 | |
| ***2. Model for ratio of antibodies to spike and antibodies to nucleoprotein for SARS-CoV-2*** | | | | 0.23 |
| Disease group | | | | |
| Control | - | - | - | |
| JIA | 0.14 | -0.01, 0.30 | 0.070 | |
| JSLE | 0.33 | 0.10, 0.58 | 0.0053 | |
| JDM | 0.50 | 0.27, 0.74 | $3.3\times10^{-5}$ | |
| Age | | | | |
| ≤13 | - | - | - | |
| >13 | -0.17 | -0.32, -0.03 | 0.019 | |
| Gender | | | | |
| Male | - | - | - | |
| Female | 0.18 | 0.06, 0.31 | 0.0035 | |
| Steroids | | | | |
| No | - | - | - | |
| Yes | 0.23 | 0.06, 0.39 | 0.0069 | |

[a]Reference categories for categorical variables indicated by “-“
